# TAD border deletion at the *Kit* locus causes tissue-specific ectopic activation of a neighboring gene

**DOI:** 10.1101/2022.12.29.522177

**Authors:** Evelyn Kabirova, Anastasiya Ryzhkova, Varvara Lukyanchikova, Anna Khabarova, Alexey Korablev, Tatyana Shnaider, Miroslav Nuriddinov, Polina Belokopytova, Galina Kontsevaya, Irina Serova, Nariman Battulin

## Abstract

Topologically associated domains (TADs) restrict promoter-enhancer interactions, thereby maintaining the spatiotemporal pattern of gene activity. However, rearrangements of the TADs boundaries do not always lead to significant changes in the activity pattern. Here, we investigated the consequences of the TAD boundaries deletion on the expression of developmentally important genes encoding tyrosine kinase receptors: *Kit, Kdr, Pdgfra*. We used genome editing in mice to delete the TADs boundaries at the *Kit* locus and characterized chromatin folding and gene expression in pure cultures of fibroblasts, mast cells, and melanocytes. We found that although *Kit* is highly active in both mast cells and melanocytes, deletion of the TAD boundary between the *Kit* and *Kdr* genes results in ectopic activation only in melanocytes. Thus, the epigenetic landscape, namely the mutual arrangement of enhancers and actively transcribing genes, is important for predicting the consequences of the TAD boundaries removal. We also found that mice without a TAD border between the *Kit* and *Kdr* genes have a phenotypic manifestation of the mutation — a lighter coloration. Thus, the data obtained shed light on the principles of interaction between the 3D chromatin organization and epigenetic marks in the regulation of gene activity.

## Introduction

Genome folding into topologically associated domains (TADs) and compartments is considered to be an important level of gene regulation, affecting the frequency of contacts between regulatory elements and gene promoters. TADs as functional units have been proposed to prevent unwanted promoter-enhancer contacts and narrow down the enhancer search space. The functional importance of TADs for gene regulation was emphasized by analyzing disease-associated mutations, disrupting their boundaries. Structural variants, crossing TAD boundaries, can provoke pathogenic gene misexpression through the rewiring of promoter-enhancer interactions (reviewed in^1,2^). Deletions have the potential to cause TADs fusion, resulting in ectopic contacts. In T cell acute lymphoblastic leukemia disruption of CTCF-associated boundary allows the proto-oncogene activation^3^. Another example is limb malformation, triggered by deletion of a TAD boundary at the *Ihh/Epha4/Pax3* locus^4^. Inversions and translocations were shown to cause TADs shuffling^4–7^. In some cases, this shuffling triggers an enhancer adoption followed by ectopic activation of the gene ^4^. In other studies, the pathogenic effect of the inversion results from the loss of expression of one of the genes with no enhancer adoption^8^. Overall, the susceptibility of a gene to an aberrant activation by an ectopic enhancer is different and partially depends on the gene’s chromatin status^6^.

Although altered TADs architecture is indeed associated with some pathogenic phenotypes^9,10^, genome wide disappearance of TADs insulation upon CTCF or cohesin depletion does not cause global transcriptional effects^11,12^. Besides, in a number of studies TADs fusion had only mild effects on gene expression pattern, as enhancers inside the newly formed TAD retained their specificity, for example in *Kcnj2–Sox9* locus^13^ and in *Shh* locus^14^. Unlike the deletion of CTCF sites, the repositioning of a TAD boundary by an inversion redirected enhancer activity and induced gene misexpression, highlighting the role of an entire TAD substructure with its epigenetic landscape in altering gene regulation^13^. Overall, how the regulatory environment inside the TAD affects the output of a boundary disruption remains an open question.

To test the effect of TAD boundary loss in different regulatory contexts, we investigated the *Kit* locus. The region comprises developmental regulators *Pdgfra, Kit* and *Kdr*, each located in the corresponding TAD, demarcated by strong domain boundaries. Mutations at this locus are often associated with malignant processes. Amplification of *Pdgfra, Kit* or *Kdr* is common among glioblastomas^15,16^. In a subset of gliomas, *Pdgfra* is aberrantly activated by an ectopic enhancer, which is associated with reduced CTCF binding^10^. *Kit* and *Pdgfra* mutations, resulting in the constitutive activation of these genes, are often found in gastrointestinal tumors^17^. Aberrant activation of *Kit* and *Pdgfra* by the oncogene-induced superenhancers is essential for leukemic cell growth and survival^18^. Finally, *Kdr* is considered one of the most critical pro-angiogenic factors in tumor angiogenesis. *Kdr* expression is upregulated in cancer cells through an autocrine loop with its ligand^19^. Thus, maintaining the integrity of the chromatin structure within the *Kit* locus appears to be important, since mis-interaction of enhancers with proto-oncogenes can activate oncogenic signals.

Here, we generated a series of genome-edited mice with deletions of boundary and intra-TAD CTCF sites and used cultured cells to track the effect of boundary perturbations. Using cell types with contrasting activity of *Pdgfra, Kit* and *Kdr* allowed us to test the role of boundary insulation in different regulatory contexts. The level of architectural changes varied from a marginal increase of inter-TAD contacts across the Pdgfra/Kit boundary, to the disrupted insulation between Kit/Kdr TADs. Loss of insulation between Kit and Kdr TADs resulted in cell type-specific ectopic gene activation. The boundary disruption was associated with a specific phenotype in mice — a lighter coat color. Collectively, our findings support the idea that the structural and functional outcome of CTCF binding sites (CBSs) deletions depends on the epigenetic landscape of neighboring TADs.

## Results

### Three adjacent TADs define 3D structure of the *Kit* locus

The *Kit* gene has been attracting the attention of researchers for a long time, as its mutations induce a clearly visible dominant white-spotting phenotype. *Kit* as well as its flanking genes *Pdgfra* and *Kdr* encode receptor tyrosine kinases, which participate in fundamental cellular processes in metazoans, such as signal transduction from growth factors and cytokines, cell proliferation and differentiation, etc. *Pdgfra, Kit* and *Kdr* genes were established from a common ancestor through a series of duplications^20^. To estimate the evolutionary conservativity of TADs in the *Kit* locus, we liftovered Hi-C contacts from several vertebrate species to the mouse genome, using C-InterSecture software^21^, since a direct comparison of Hi-C maps from different species is inaccurate due to different genome sizes (Supplementary Fig.1a). Resulting interaction maps demonstrate a high similarity of chromosome architecture across the *Kit* locus in human, mice, dog, chicken^22^ and african clawed frog^23^ (Supplementary Fig.1b).

Each of the three genes features a distinct tissue-specific expression pattern. *Pdgfra-positive* cells can be found in a wide array of mesenchymal tissues^24–26^. *Kit* is expressed among several tissues, particularly in mast cells and interstitial cells of Cajal, being among the most expressed genes in mast cells (Fig.1a). *Kdr* is an endothelial-specific receptor and its expression is almost exclusively restricted to the blood vessels^27,28^. In order to evaluate the effect of CTCF sites deletion in different regulatory landscapes, we focused on three cell types with contrast expression of the *Pdgfra, Kit, Kdr* genes: fibroblasts *(Pdgfra* is expressed, *Kit* is inactive), mast cells (*Kit* is among top 0.3% of the expressed genes, *Kdr* is inactive), and melanocytes (*Kit* is among top 3% of the most expressed expressed genes, *Kdr* is weakly expressed) (Fig.1a). While the use of animal tissues results in an averaged picture due to the presence of many cell types in the sample, using a pure population of cultured cells allowed us to obtain more precise genomic data (Supplementary Fig.2).

**Fig. 1.**
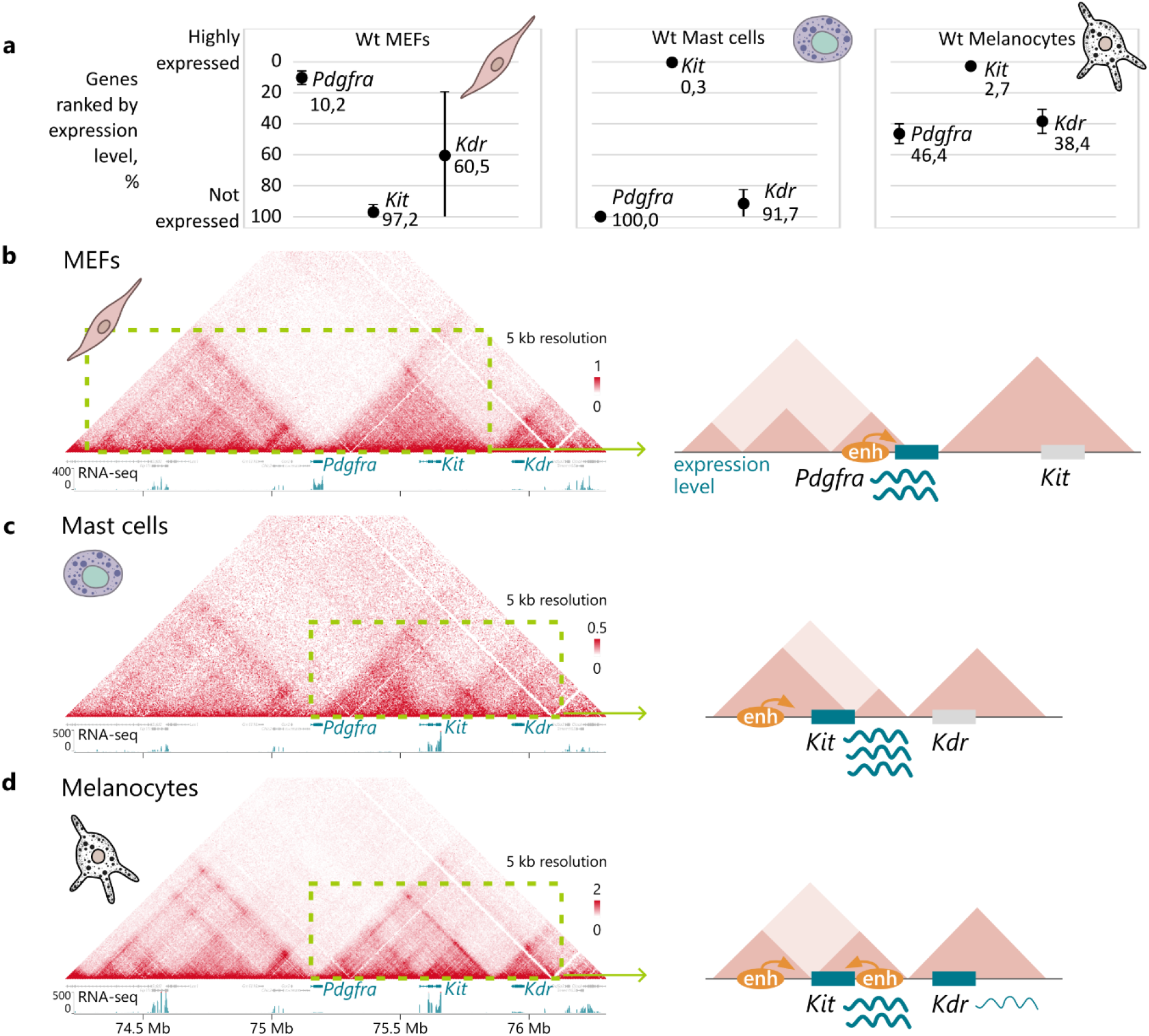
Spatial and regulatory genome characteristics of the Kit locus. **a** The genes *Pdgfra, Kit, Kdr* demonstrate contrasting expression patterns in the chosen cell types, as evidenced by the RNA-seq data analysis. Error bars represent the standard deviation, n=3; Tissue-specific TAD organization of the Kit locus in mouse embryonic fibroblasts (MEFs) (**b**), mast cells (**c**) and melanocytes (**d**) corresponds to different enhancer localization and expression level (schemes on the right). Colour bars reflect the interaction counts.

**Fig. 2.**
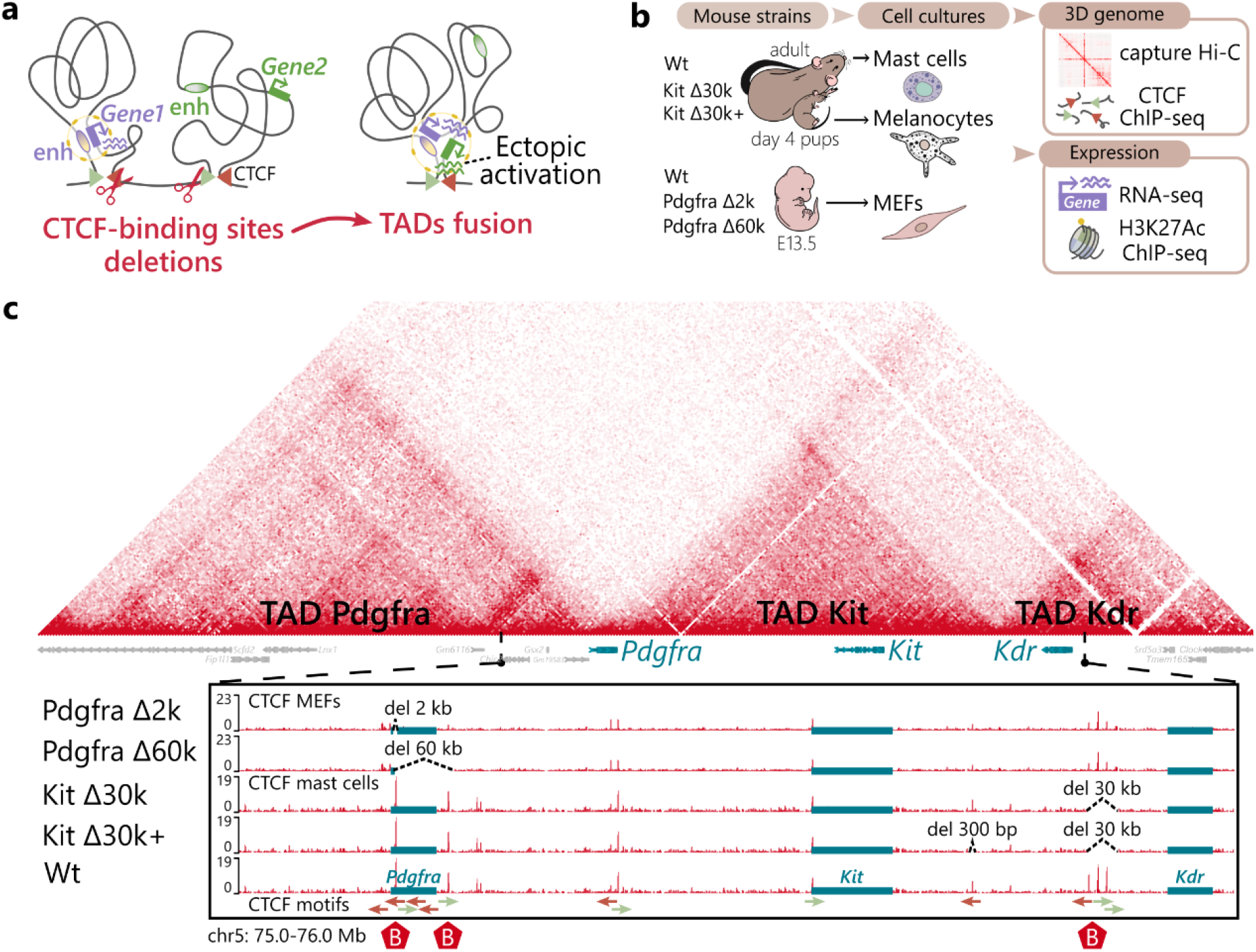
Experimental design and generated mutations. **a** Schematic of enhancer interactions rewiring induced by CTCF-binding sites deletion; **b** Overview of the experiment: generation of mutant mouse strains with TAD boundary deletion, obtaining cell cultures (mast cells, melanocytes and mouse embryonic fibroblasts, or MEFs) and analysis of 3D genome organization and gene expression; **c** Schematic of CTCF binding sites deletions in mutant mouse strains, illustrated on cHi-C from MEFs and CTCF ChIP-seq from MEFs and mast cells. Arrows (red-green) indicate CTCF binding sites orientation.

To characterize the TADs organization in mouse embryonic fibroblasts (MEFs), mast cells and melanocytes we performed capture Hi-C (cHi-C). Our data confirmed that the region is organized into three adjacent TADs with *Pdgfra, Kit* and *Kdr* each located in a separate corresponding TAD (Fig.1b-d). Notably, the *Pdgfra* gene body overlaps an extended region between Pdgfra and Kit TADs, while its promoter is located immediately upstream of the Pdgfra TAD boundary. It is crucial for enhancers to physically interact with target promoters, therefore the *Pdgfra* localization suggests that the TAD border is robust. As to *Kit* and *Kdr*, their coding regions are located within the corresponding TADs. Importantly, two self-interacting regions make up the Kit TAD in mast cells and melanocytes. We noted that in mast cells the insulation between these regions coincides with the *Kit* terminator, while in melanocytes it corresponds to the *Kit* promoter.

According to our H3K27Ac ChIP-seq data (schemes at Fig.1 and Fig.3-4), *Pdgfra* and *Kit* enhancers localize upstream of the genes in MEFs and mast cells, respectively. In melanocytes, by contrast, the H3K27ac-rich enhancer region is located downstream. Different localization of *Kit* enhancers in mast cells and melanocytes probably accounts for the position of an intra-TAD insulation. Thus, in mast cells *Kit* with its enhancers is spatially confined within the left half of the Kit TAD, and in melanocytes within the right half.

**Fig. 3.**
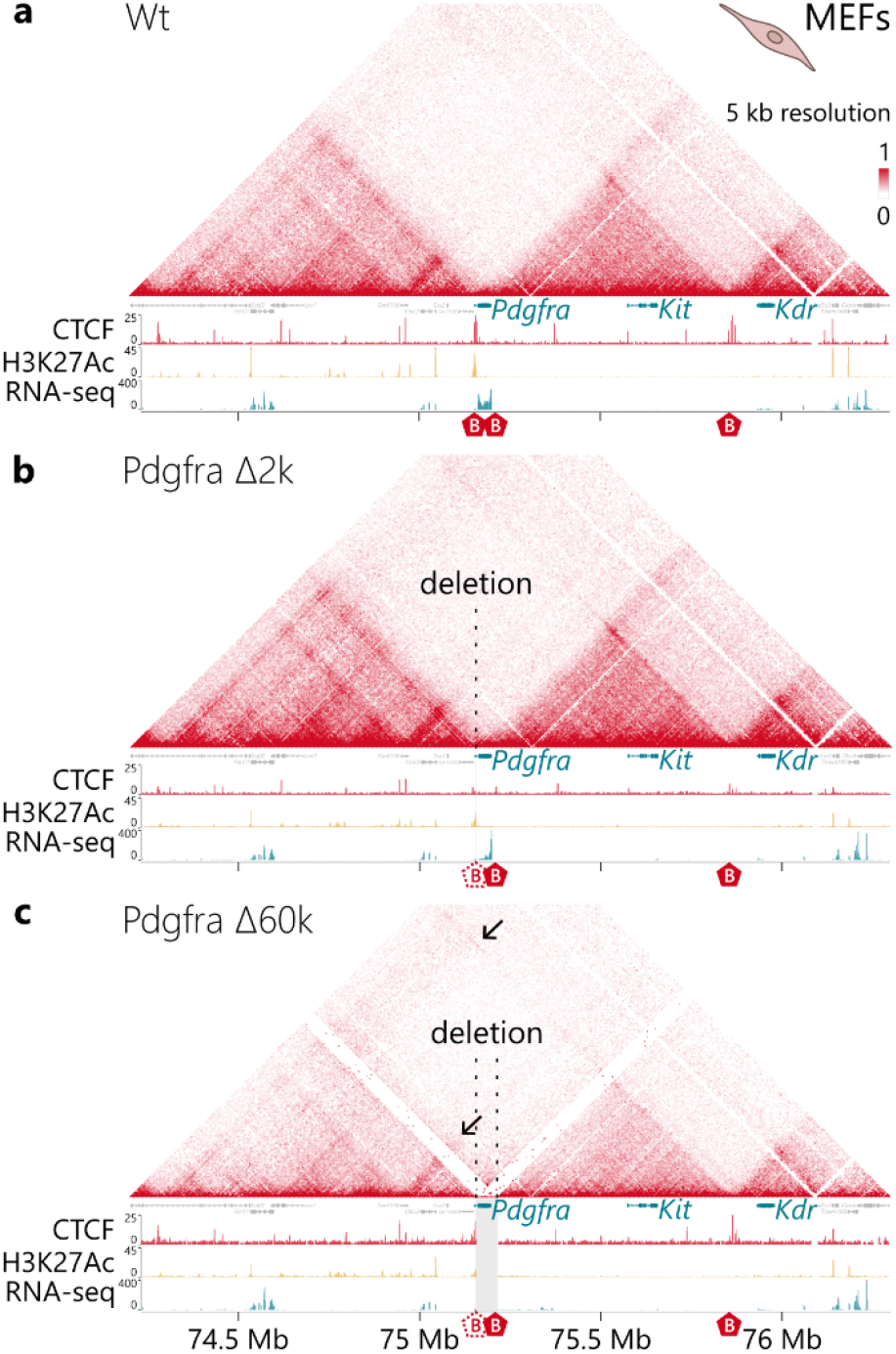
3D chromatin structure of the *Kit* locus in MEFs. CBSs deletions Pdgfra Δ2k and Pdgfra Δ60k were insufficient to disrupt the Pdgfra/Kit TADs insulation and change the regulatory pattern in mouse embryonic fibroblasts (MEFs). Hi-C heatmaps, RNA-seq and ChIP-Seq signals across the *Kit* locus in wild-type (**a**), Pdgfra Δ2k mutant (**b**) and Pdgfra Δ60k mutant (**c**) MEFs. Ectopic contacts in Pdgfra Δ60k are indicated by arrows.

**Fig. 4.**
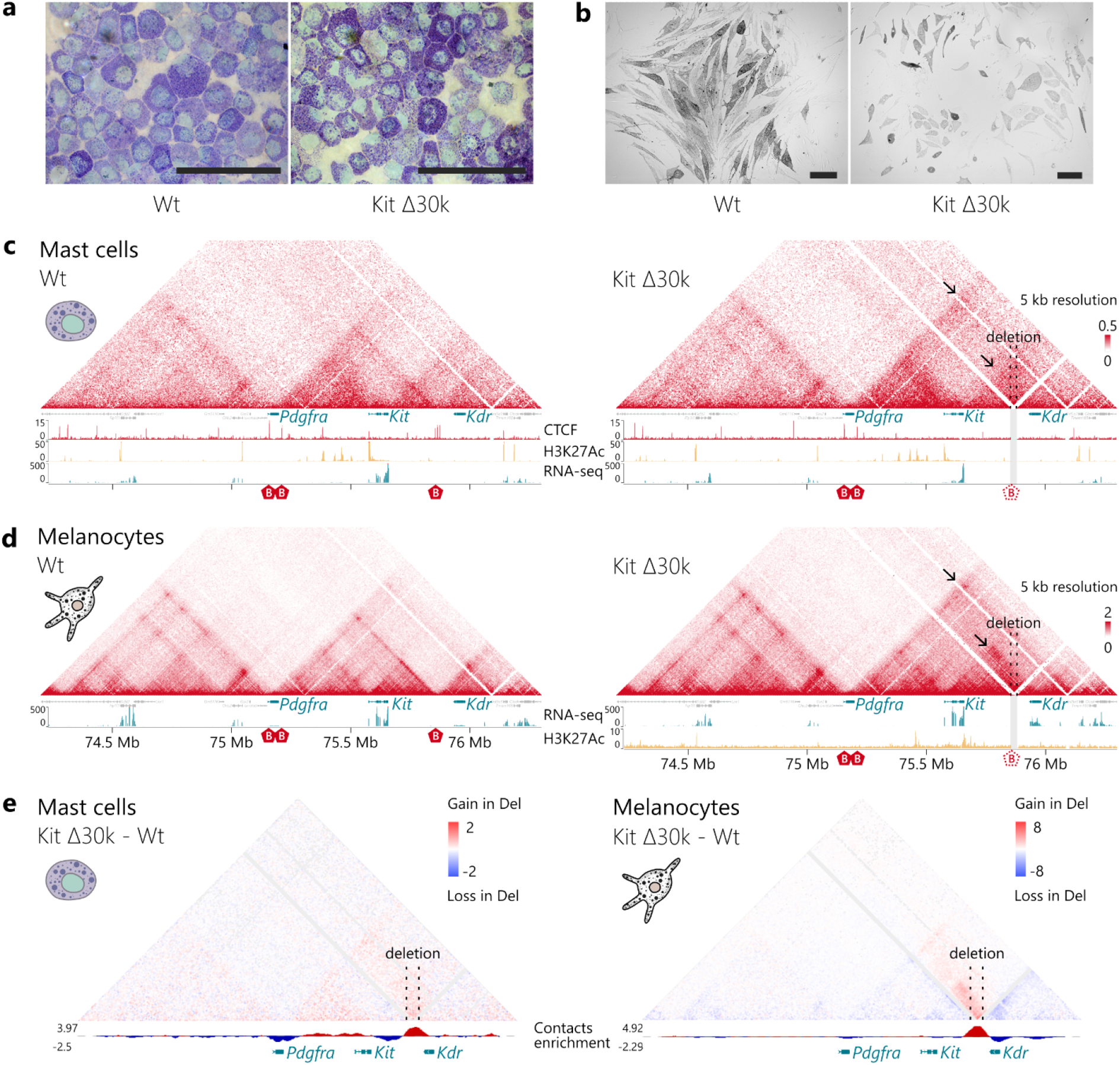
3D genome organization of the *Kit* locus in mast cells and melanocytes. **a** Cytospin preparations of wild-type (left) and Kit Δ30k (right) mast cells after 4 weeks in culture. Histamine containing granules could be detected by toluidine blue staining. Scale bar: 100 um; **b** Primary wild-type (left) and Kit Δ30k (right) melanocytes after 2.5 weeks in culture. Scale bar: 100 um. **c**-**d** cHi-C heatmaps, RNA-seq and ChIP-Seq signals across the *Kit* locus in wild-type (left) and KitΔ30k (right) mast cells (**c**) and melanocytes (**d**); ectopic inter-TAD interactions are indicated by arrows; **e** Subtraction maps showing changes in contact frequencies in mutant cells relative to the wild-type in mast cells (left) and melanocytes (right). The tracks below the subtraction maps show the estimated contact enrichment, indicating loss of insulation in the mutant cells. Removal of the Kit/Kdr boundary resulted in loss of inter-TAD insulation and extensive interactions across the TADs boundary.

The spatial architecture and the regulatory pattern of the *Pdgfra/Kit/Kdr* locus demonstrate tissue-specific characteristics. Comparing the effects of CBSs deletions in different cell types allowed us to analyze how the deletions influence the TADs integrity in different regulatory landscapes.

### Generated mouse strains carrying boundary CBSs deletions

We expected the removal of the boundaries to result in TADs fusion, permitting enhancer hijacking and aberrantly elevating gene expression (Fig.2a). To assess whether TADs restrict an enhancer influence to its target promoter, we generated a series of mouse strains carrying different deletions of the key CBSs either at the Pdgfra/Kit or Kit/Kdr TADs boundaries, obtained primary cell cultures from the mice and performed an analysis of 3D genome and gene expression (Fig.2b). For a comprehensive view on the CTCF binding motifs at the *Kit* locus see Supplementary Fig.3.

The Pdgfra/Kit boundary area comprises boundaries of two adjacent TADs separated by a 50 kb region with the *Pdgfra* gene body located in that space. Our ChIP-seq data and further consensus motif identification revealed that the Pdgfra TAD border is characterized by three strong CBSs in reverse orientation, located within the *Pdgfra* gene body (Fig.2c). The **Pdgfra Δ2k** strain carries a 2 kb deletion of the three CTCF peaks at the Pdgfra TAD border. But a CTCF peak at the Kit TAD boundary remained intact. Thus, the Pdgfra Δ2k strain represents a case with a partial disruption of the border’s structure, which allowed us to check whether the three CTCF sites at the Pdgfra side of the boundary region are required for the Pdgfra/Kit TADs integrity. Still, *Pdgfra* enhancers remained isolated from the *Kit* promoter by the Kit TAD border. Kit TAD boundary is characterized by a single forward-oriented CBS, which was further deleted in the **Pdgfra Δ60k** strain. Thus, Pdgfra Δ60k strain carries a 60 kb deletion spanning the whole Pdgfra/Kit TAD boundary region, and therefore removing the *Pdgfra* coding sequence, leaving only 3 out of 22 exons and a promoter. Even though initially we intended to simultaneously introduce two separate deletions at the Pdgfra and Kit sides of the boundary region, we considered the Pdgfra Δ60k strain valid and sufficient for the research purpose despite the *Pdgfra* knockout, since the promoter and enhancers of the *Pdgfra* gene remained intact.

The Kit/Kdr boundary is a joint between two adjacent TADs which contains three clustered CBSs, with two sites in reverse and one in forward orientation (Fig.2c). The **Kit Δ30k** mouse strain carries a 30 kb deletion of the three CBSs at the Kit/Kdr border. Analyzing the first Hi-C heatmaps performed on the Kit Δ30k cell cultures, we noticed a strong CTCF ChIP-seq signal within the Kit TAD, located ~116 kb away from the boundary. To minimize any potential impact of this site in maintaining the TAD’s structure, we generated a **Kit Δ30k+** strain, carrying an additional 300 bp deletion.

Performed deletions only mildly affected the mice phenotype, except the Pdgfra Δ60k. Since *Pdgfra* is crucial for the early development, its deletion in homozygous state is lethal for embryos by 15 embryonic day (E15), so the Pdgfra Δ60k strain was maintained in a heterozygous state. Nonetheless, we were able to obtain MEFs cultures from the E13.5 homozygous embryos. For Kit Δ30k, we detected the color of mice changing from black to brown relative to the wild type. It is an interesting feature that we further discuss in the section with analysis of Kit Δ30k mutation effects.

The generation of four mutant mouse strains, carrying different deletions of the key CBSs either at the Pdgfra/Kit (Pdgfra Δ2k, Pdgfra Δ60k) or Kit/Kdr (Kit Δ30k, Kit Δ30k+) boundary, allowed us to perform a comprehensive research of TADs disruption influence on the locus.

### Pdgfra/Kit boundary CBSs deletions were insufficient to change the regulatory and architectural patterns

To elicit whether CBSs deletions at the Pdgfra/Kit boundary change regulatory and expression patterns of the genes, we analyzed MEFs obtained from E13.5 embryos. *Pdgfra* and *Kit* show a contrasting expression in MEFs, i.e. *Pdgfra* is actively transcribed, while *Kit* is not transcribed (Fig.1a). In agreement with our hypothesis, we expected *Pdgfra* enhancers to induce *Kit* transcription as a result of TADs fusion in an absence of boundary CBSs.

To assess the spatial architecture of the locus, we performed CTCF ChIP-seq and cHi-C on MEFs from the Pdgfra Δ2k and Pdgfra Δ60k mice (Fig.3). The Pdgfra Δ2k deletion, affecting the 2 kb region at the Pdgfra TAD border, had no detectable effect on 3D contacts. This could be explained by the remaining CBSs at the Kit side of the Pdgfra/Kit boundary. By contrast, in Pdgfra Δ60k deletion, affecting a larger region spanning both sides of the Pdgfra/Kit border, the effect was more distinct. The observed interaction profiles were confirmed by the evaluation of inter-TAD contacts enrichment in mutant cells relative to the wild-type (Supplementary Fig.4 and Fig.5a,b). These results are in good agreement with the reported robustness of TAD borders with larger deletions of boundary CBSs more likely to trigger the TADs fusion^14,29^.

Pdgfra Δ60k deletion causes a visible stripe of ectopic contacts to establish between Pdgfra and Kit TADs despite a strong insulation between the two TADs still being observed. In the Pdgfra Δ60k mutants, the *Pdgfra* promoter is retained, containing a small CTCF peak, located at a 4kb distance within the Pdgfra TAD (Fig.3c). The persistence of this CTCF site may account for the insulation remaining between the TADs. Upon the deletion of boundary CBSs, a TAD boundary can shift towards the position of the nearest intra-TAD CTCF site, as was shown for the HoxA cluster during motor neuron differentiation^30^. In the present study, we might see a similar compensating mechanism that ensures the formation of the 3D architecture. Thus, the Pdgfra and Kit TADs remained largely separate despite the successful deletions of the boundary CBSs.

We further asked if the mutations changed the expression pattern. To explore the transcriptome, we performed RNA-seq of the mutant and wild-type MEFs. In Pdgfra Δ60k, an intergenic region between *Pdgfra* and *Kit* is transcribed, starting from the intact *Pdgfra* promoter and continuing to the nearest termination site. RNA-seq data revealed no increase in *Kit* transcriptional activity in any of the mutant MEFs lines. Consistent with our transcriptome analysis, an H3K27ac-marked enhancer region of the *Pdgfra/Kit* locus remained similar between the mutant and wild-type MEFs lines (Fig.3).

The obtained data shows that neither local CBSs deletion Pdgfra Δ2k, nor the larger Pdgfra Δ60k disrupted the Pdgfra/Kit TADs insulation. Therefore, the CBSs deletions were insufficient to change the regulatory pattern and to activate *Kit* expression in fibroblasts.

### Kit/Kdr boundary CBSs deletions result in the TADs fusion

Similarly, we deleted three CTCF sites, which delineate the Kit/Kdr border, and monitored the effects of the deletion on two cell types where *Kit* is actively transcribed — mast cells and melanocytes. *Kdr* is normally inactive in mast cells and is marginally expressed in melanocytes (Fig.1a).

Mast cells were differentiated *in vitro* from bone marrow-derived progenitors. The resulting cell population was highly homogeneous and displayed the typical morphological features, with heparin and histamine containing granules that can be detected by toluidine blue staining (Fig.4a). We also used flow cytometry to verify the purity of the population (Supplementary Fig.2b).

Contact maps from the wild-type mast cells denoted a nested substructure of the Kit TAD, with two self-interacting regions. *Kit* with its upstream enhancers are located within the right half of the TAD and thus are insulated from the left half (Fig.1c). Kit Δ30k deletion disrupted the Kit/Kdr TAD boundary insulation in mast cells and led to a noticeable gain of inter-TAD contacts (Fig.4c). A similar pattern was found for Kit Δ30k+ (Supplementary Fig.4). Looping interactions connecting the outer boundaries of Kit and Kdr TADs were also established. However, we noted that contacts rewiring mainly involved the right half of the Kit TAD, which merged with the Kdr TAD. The insulation inside the Kit TAD was not affected by the deletion, thus, a self-interacting region, located at the right side, remained isolated. Kit Δ30k+ deletion, where an additional CBS within the Kit TAD was removed, had a similar effect on contact frequency and did not lead to a discernible contacts rewiring compared to Kit Δ30k (Supplementary Fig.4). Thus, despite the TAD boundary disruption in mast cells, active *Kit* enhancers remained insulated from *Kdr*.

We examined the CTCF binding in wild-type and mutant mast cells (Fig.4c). Our ChIP-seq analysis did not reveal any CTCF peaks at the boundary between two self-interacting regions inside the Kit TAD. We also did not observe any novel CTCF peaks in mutant cells, similar to the Pdgfra/Kit boundary deletion. Hence, *Kit* enhancers are possibly restrained from the downstream genes by different mechanisms, probably through the *Kit* transcriptional activity.

We then compared the effects of the boundary CBSs deletion in mast cells and melanocytes. Melanoblasts from the skin of 4-day-old mice were differentiated *in vitro*. Resulting cell population, consisting of granulated melanocytes (Fig.4b and Supplementary Fig.5), was assessed by flow cytometry (Supplementary Fig.2b). The inner structure of the Kit TAD in melanocytes is determined by the downstream localization of *Kit* enhancers, together with *Kit* forming a self-interacting region in the left half of the TAD (Fig.1d). Kit Δ30k deletion in melanocytes resulted in extensive rewiring of chromatin contacts. Our contact maps revealed loss of insulation at the Kit/Kdr boundary and concurrent merging of neighboring TADs (Fig.4d). Similarly with mast cells, gain of interactions was especially noticeable between the left half of the Kit TAD and *Kdr* region. But in melanocytes the boundary deletion induced the fusion of *Kit* and its enhancers with *Kdr* region into a new substructure, enabling *Kit* enhancers to interact with the *Kdr* promoter.

Thus, in both cell types, removal of boundary CTCFs resulted in loss of inter-TAD insulation and extensive interactions across the TADs boundary. At the same time, the insulation between two self-interacting regions within the Kit TAD was preserved. Our results imply that this insulation is provided by the epigenetic environment, in particular, by the relative position of enhancers and actively transcribed *Kit* within the TAD.

### Eliminating the contribution of distance decrease to the Hi-C interaction profiles in mutant cells

In addition to the epigenetic factors, the distribution of genomic contacts is determined by a distance between loci. Since some of the deletions spanned large genomic regions (30 and 60 kb), we sought to eliminate the contribution of distance decrease to the rewiring of chromatin interactions detected in cHi-c maps from mutant cells. For this purpose, we generated wild type Hi-C maps with simulated deletions (Supplementary Fig.6). To do this, we remapped our wild type Hi-C contacts on a custom mm10 genome containing the deletion. The remaining contacts were recalculated according to a P(s) decay. We used the same C-InterSecture algorithm to remap Hi-C contacts from different species to the mouse genome for the estimation of evolution conservativity of 3D structure of the *Kit* locus. The subtraction maps of mutant cells and wild-type controls display major chromatin rewiring, indicating that the establishment of inter-TAD contacts was due to the removal of CTCF sites and not to the reduction of genomic distance (Fig.4e and Supplementary Fig.7). Using this approach allowed us to unbiasedly assess the effect of boundary CTCF sites deletion.

### The Kit/Kdr TADs fusion provides tissue-specific ectopic gene activation

We then asked whether the reorganization of contacts observed in the Kit Δ30k and Kit Δ30k+ mutant mice was functional and could trigger a transcriptional outcome.

In mast cells, despite the boundary disruption, the H3K27ac-marked enhancer region did not gain any substantial ectopic contacts with the downstream promoters (Fig.4c). Consistent with this, RNAseq analysis in mutant mast cells revealed no transcriptional changes in the locus or among the adjacent genes (+/- 5Mb).

Remarkably, the situation was different in melanocytes. *Kdr* is normally expressed in melanocytes, although at a low level. In Kit Δ30k cells, we detected a 6.7-fold increase in *Kdr* expression level compared to the wild-type data (Fig.5a). Ectopic transcriptional activation observed in mutant melanocytes is in agreement with the redirection of the promoter-enhancer contacts.

**Fig. 5.**
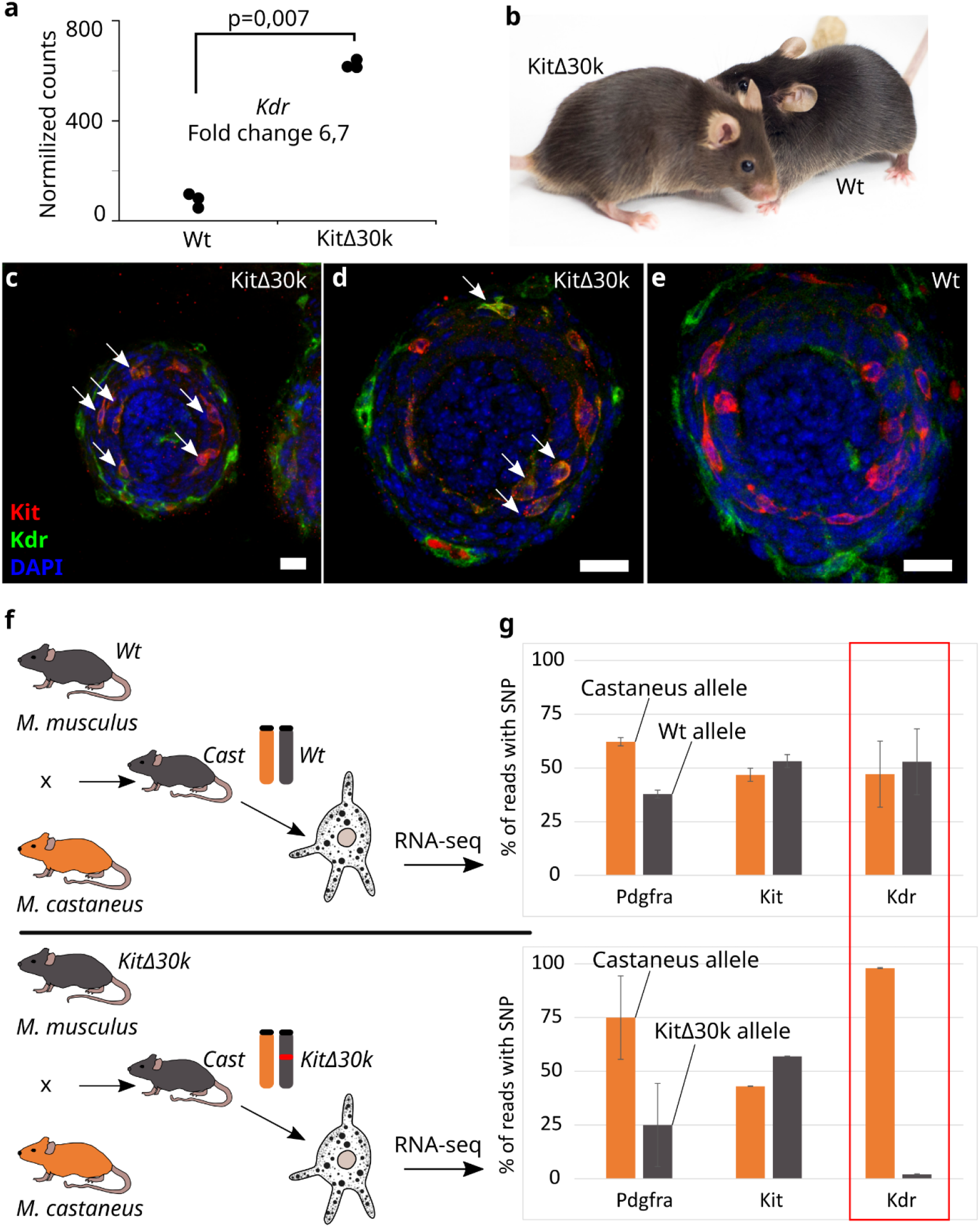
Kit/Kdr TADs fusion induces *Kdr* activation in melanocytes and phenotypic changes. **a** A plot of normalized RNA-seq reads counts in wild-type and Kit Δ30k melanocytes indicates a 6.7-fold increase of *Kdr* transcription. Black dots represent individual replicates. p-value was estimated with DESeq2, using the Wald test; **b** Adult mice with Kit Δ30k deletion had brownish fur color (left) compared to the black wild-type animals (right); **c,d** Vibrissae hair bulge section of KitΔ30 E15.5 embryo. Arrows — double *Kit* (red) and *Kdr* (green) positive cells; **e** Vibrissae hair bulge section of wild-type E15.5 embryo. Scale bar: 20 um; **f** Crossing scheme used to obtain *Mus musculus* x *Mus castaneus* hybrids; **g** Percentage of reads with *M. castaneus-specific* SNPs was scored for the evaluation of allelic activity of *Pdgfra, Kit* and *Kdr* in hybrid melanocytes. Number of SNPs within the genes: *Pdgfra —* 36, *Kit* — 16, *Kdr —* 19.

We next set out to detect an aberrant *Kdr* expression in melanocytes *in vivo*. We performed immunostaining in E13.5 and E15.5 embryos, focusing on several areas on the cryosections: somites, yolk sac, skin areas on the head and back (Supplementary Fig.8), eyes, and vibrissae hair bulge (Supplementary Fig.9). We looked for double-positive cells, expressing both *Kit* and *Kdr. Kit* expression is detected in a large number of cell types, while *Kdr* is mostly restricted to the heart and vascular system. In Kit Δ30k embryos double-positive cells were detected in the vibrissae hair follicles at E15.5 (arrows at Fig.5c-e). Location of these cells suggests that these are melanocytes^31^.

We also found single double-positive cells, located alongside blood vessels in the skin of E13.5 embryos (Supplementary Fig.9i-p). We cannot accurately identify these cells, as we have not performed additional surface markers staining. These might be *Kit*-positive cells of the hematopoietic lineage^32^. Notably, we did not detect *Kit* activation in *Kdr*-expressing endothelial cells, at least immunohistochemically. It can be assumed that in endotheliocytes, Kit Δ30k deletion did not cause the *Kdr* enhancer hijacking.

We noted that Kit Δ30k deletion was associated with a specific phenotype in adult mice. Mutant mice had brownish fur color, compared to the black wild-type animals (Fig.5b). We performed a manual counting of melanocytes on immunostained skin sections from 4-day-old pups aiming to explain the specific phenotype of the paler Kit Δ30k mice (Supplementary Fig.10). However, the analysis did not reveal significant differences between two genotypes; hence, the mechanism of mutation-associated phenotypic manifestation is still in question.

Thus, the deletions of the boundary CBSs at the Kit/Kdr TADs border induced tissue-specific effects. For mast cells, despite the weakened insulation between two TADs, the *Kit* enhancer region did not form new contacts with downstream promoters and *Kdr* remained inactive. In melanocytes, TAD boundary deletion induced the rewiring of *Kit* enhancers, which triggered *Kdr* misexpression and phenotype changes. We propose that the individual epigenetic characteristics of the locus, e.g., gene activity and enhancer localisation within the TAD, determine the tissue-specific response to a boundary deletion.

### *Mus musculus* x *Mus castaneus* breeding further confirmed ectopic *Kdr* activation to result from boundary CBSs deletions

During the melanocytes *in vitro* differentiation we noticed that the cells obtained from the Kit Δ30k and Wt mice looked slightly different. Kit Δ30k melanocytes were paler and did not accumulate melanosomes as actively as Wt cells (Fig.4b). Since the culturing process takes 4.5-7 weeks, it can be assumed that the cells were under an influence of confounding effects (cultivation conditions, trans-acting factors, differentiation state, etc.). Therefore, it is possible that increased *Kdr* expression does not reflect the difference in the activity of genes on the chromosome with and without deletion, but is only a consequence of different transcriptional status of cells. To test this hypothesis, we crossed Kit Δ30k mice with *Mus castaneus* mice (Fig.5f). Since almost all genes carry SNPs that distinguish *M. musculus* from *M. castaneus*, we were able to evaluate the activity of the gene in comparison with the reference (castaneus) allele. Since both alleles are in the same cell, this assessment does not depend on transcriptional status of cells. We isolated primary cells from the epidermis of the hybrids, differentiated them, performed RNA-seq in melanocytes, and analyzed the proportion of reads carrying SNPs from one of the two species. We have shown that almost all (98%) *Kdr* transcripts in hybrid melanocytes originate from the chromosome carrying the Kit Δ30k deletion (Fig.5g). The overall level of *Kdr* expression was significantly lower in the control experiment with *M. castaneus* x *M. musculus* Wt hybrids, with both alleles being equally active. Thus, ectopic activation of the *Kdr* gene in melanocytes is caused by a deletion of the TAD border between the *Kit* and *Kdr* genes.

## Discussion

We found that TADs contribution to gene regulation depends on tissue-specific features at the *Kit* locus. The TAD border deletion caused fusion of the Kit and Kdr TADs in mast cells and melanocytes. Whereas an ectopic activation of *Kdr* occurred only in melanocytes. Several studies demonstrated that whether border CBSs deletion causes an ectopic gene activation largely depends on the locus, since the deletions provide inconsistent results (Supplementary Table 1). For the first time we provide comparison of the same locus at different regulatory landscapes in several pure primary cell cultures. Our results indicate that TADs may unite different regulatory features, thus their function is not limited to an integrity of CBSs border, as was previously assumed.

### *Kit* transcription defines tissue-specific characteristics of the TAD

We analyzed the effect of TADs disruptions in primary cell cultures, as opposed to an averaged data from tissues largely used before^4,13,29,33,34^, which allowed us to obtain high-quality genomic data. The data provided a view on chromatin interactions at different regulatory landscapes of the *Kit* locus. Three adjacent TADs encompass the locus in MEFs, but in mast cells and melanocytes the Kit TAD is additionally divided into two.

Since *Kit* is inactive and an insulation is absent in MEFs, we presume transcription to provide the insulation. It is in agreement with the study revealing that transcription modulates chromosome folding during mouse thymocyte maturation with single activated genes forming an insulated region within TADs^35^. Models of cohesin loading and transcription at different states suggested RNA polymerase II (RNAP) to act as a barrier to loop extrusion^36^. Transcribing RNAP may disrupt chromatin interactions by displacing cohesin from CTCF sites^37,38^ relocating it over long distances along DNA^39^. Moreover, upon depletion of RNAP many enhancer-promoter loops are lost and new loops are established at the CTCF sites, which were restricted by RNAP^40^.

Notably, the border of an insulated region coincides with a CTCF site at the *Kit* promoter in melanocytes, whereas it shifts toward the *Kit* terminator in mast cells. Though *Kit* is expressed in both cell types, the expression level in mast cells is 10 times higher. High level of transcription is assumed to antagonize CTCF-cohesin loop extrusion^41^. Cohesin was revealed to accumulate at 3’-end of highly active genes in an absence of CBS or an efficient cohesin release by WAPL^39,42^.

In addition, transcription can participate in the formation of nested structures within TADs, which are established independently of CTCF-cohesin loop extrusion mechanism^43^. It could explain why the insulated region within the Kit TAD in mast cells is not as distinct as in melanocytes, where it is formed in a CTCF-dependent manner.

Thus, a high level of *Kit* expression could influence tissue-specific features of the spatial genome organization at the locus.

### Deletion of a border between two TADs causes their fusion

We detected that removal of border CBSs resulted in TADs fusion, though architectural changes varied from a marginal increase of inter-TAD contacts across the Pdgfra/Kit border to the full contacts enrichment across the Kit/Kdr border.

We assume the CTCF site at the *Pdgfra* promoter, which remained intact after the deletions, to account for an incomplete loss of an insulation between Pdgfra and Kit TADs. It is in agreement with the fact that CBSs might be dispensable to maintain a TAD border ^14,44^. Though an insulating strength of an individual CBS is not correlated with its CTCF occupancy^45^, which corresponds to our CTCF ChIP-seq data showing similar CTCF occupancy of the remaining site at the *Pdgfra* promoter between mutant and wild-type MEFs. Another potential explanation is that transcription from the *Pdgfra* promoter located at the TAD border — transcripts being the gene at the Pdgfra Δ2k and an intergenic region at the Pdgfra Δ60k — could strengthen the insulation. It was proposed that transcribed genic and non-genic boundaries are associated with stronger insulation due to boundary RNAs facilitating CTCF recruitment^46^.

Thus, Pdgfra and Kit TADs remained largely separate. Since an enhancer influence on the promoter is constrained by TADs borders^47^, a few newly established spatial interactions were insufficient for *Kit* activation in MEFs. There is another mouse strain carrying the deletion of *Pdgfra* gene — *Patch* mutation. Interestingly, it causes an ectopic *Kit* activation in somites and lateral mesenchyme and exhibits a distinct pigment phenotype — white abdomen ^48–50^. An analysis of changes in genome and spatial organization of the *Patch* mutant mice is yet to be performed. We suspect that the mutation covers a larger portion of the genome, which allows it to result in TADs fusion and consequently enhancer hijacking.

A limitation of our study is that for this border we provide an analysis of only one cell type. As an example, glia has the same expression profile of *Pdgfra* and *Kit* and its comparison to MEFs could be interesting. We were able to analyze the TAD border disruption only on *Pdgfra* knockout mice, which is lethal in homozygous state. To disrupt the borders simultaneously and locally (to leave *Pdgfra* intact) is a challenging task, though there is an option of generating two deletions successively, but it requires a much longer process.

As for the Kit/Kdr border, loss of insulation resulted in fusion of the TAD part downstream of the *Kit* promoter with the Kdr TAD according to the insulation pattern within the Kit TAD in mast cells or melanocytes. We emphasize that it is important to eliminate an impact of decrease in genomic distance due to deletions on the rewiring of chromatin interactions.

### TAD disruption results in *Kdr* activation in melanocytes, but not in mast cells

How TADs integrity affects gene regulation is a matter of great interest, given TADs are assumed to establish enhancer-promoter interactions.

Our transcriptome analysis of cells with disrupted TADs at the *Kit* locus revealed an ectopic activation of *Kdr* in mutant melanocytes, whereas *Kdr* remained silent in mutant mast cells. We additionally confirmed *Kdr* activation in melanocytes to result from the Kit Δ30k deletion by breeding Kit Δ30k *M. musculus* with wild-type *M. castaneus*, showing that the vast majority of the *Kdr* transcripts (98%) originated from a *M. musculus* allele. The observed difference corresponds to the 3D architecture of the locus upon Kit Δ30k deletion. Interestingly, even a ten times larger deletion at the Kit/Kdr border (300 kb against 30 kb in Kit Δ30k)^51^ does not cause *Kdr* activation in mast cells, despite a serious decrease of genomic distance between *Kit* enhancers and *Kdr* promoter (Supplementary Fig.11). Of note, we did not detect a reverse situation of *Kit* activation by active enhancers of *Kdr* in endothelial cells. Thus, cell type-specific features of the locus, such as localization of H3K27Ac-marked active enhancers, underlie the TADs function, at least in case of the *Kit* locus (Supplementary Fig.12).

Cell type-specific effect of TADs disruption on gene expression was also demonstrated in comparisons of induced pluripotent stem cells (iPSCs) and iPSC-derived cardiomyocytes^52^, ESCs and motor neurons^30^. In addition, strength of TADs border may vary between cell types, as was shown for mESCs and MEFs^44^. Though given a generally more open chromatin state of stem cells relative to differentiated cells^53^, the advantage of our study is providing evidence of the effects in primary cell cultures from animals with deletions.

Studies concerning the influence of TADs disruption on expression suggest that regulation within TADs may largely depend on other factors rather than on the integrity of a TAD border only (Supplementary Table 1). We provide evidence of cell-type specific features of the *Kit* locus to affect a TAD’s role insulating an enhancer influence on a non-target promoter.

### Phenotypic consequences of the Kit TAD disruption

In our study we observed a mice phenotype change as a result of the CBSs deletions at the border between Kit and Kdr TADs: the fur of the Kit Δ30k mice were lighter relative to the wild-type mice. It is well-known that mutations in *Kit* are implicated in the white-spotting phenotype in mice^54–56^, rabbits^57,58^, cats^59^, dogs^60^, horses^61^ and other animals. The mechanism for this phenotype is an impaired melanocyte development and migration due to the defective Kit receptor signaling. Unfortunately, we were unable to detect the significant difference in melanin accumulation or melanocytes migration between mutant and wild-type mice, possibly because such a relatively small change in coloration requires a very small change in melanocyte behavior. However, it is obvious that coloration is an essential object for natural selection, so even such an almost invisible change in the laboratory could be eliminated in natural conditions.

Mutations leading to TADs fusion are under negative selection^62^, thus, the evolutionary conservatism of TADs may be a consequence of their importance for maintaining a certain pattern of gene activity. Whereas a change in the structure of TADs can lead to the emergence of a new pattern of gene activity, which may be an evolutionary innovation. This was remarkably demonstrated by the example of moles^63^. One of the adaptations of moles to an underground lifestyle is to increase the concentration of testosterone in females. An aberrant pattern of *Fgf9* gene activity ensures the tolerance of female oogenesis to the male hormone. To change *Fgf9* activity, a chromosomal rearrangement, affecting the distal part of the *Fgf9* TAD, occurred in moles. Given TADs role in insulating a regulatory pattern, they ensure an integration of a newly established gene or a change in regulation to occur without huge alterations in neighborhood^64^. Thus, the regulatory context provided by TADs is an important object for the evolution of organisms.

Since TADs are conservative between cell types of close species^65,66^ and genes are differentially expressed between tissues, the mechanism underlying the regulation within TADs, observed in the study, ought to be wide-spread.

### Perspective

We conclude that different effects of TAD border deletions may largely depend on the characteristics of an individual locus in an exact cell-type specific context. We revealed that an epigenetic landscape in the *Kit* locus, especially an active transcription and an enhancer localization, accounts for a different insulating role of the *Kit* TAD in mast cells and melanocytes. We suggest further research to focus on how and in what ratio these features impact the TADs insulation role.

## Methods

### Animals

Mouse Kit Δ30k line was established from a chimeric animal generated after mouse embryonic stem cells (mESCs) injection as described earlier^67^. Briefly, mESCs were transfected with CRISPR/Cas9 plasmids aimed at a border region between Kit and Kdr TADs (see gRNA in Supplementary Table 2). With PCR genotyping we selected a homozygous clone with Kit Δ30k deletion (see primers in Supplementary Table 2). Modified mESCs were injected into recipient blastocysts. Chimeric founders (F0) were crossed with wild-type (Wt) C57BL/6J animals to generate heterozygous animals (F1). After 8 rounds of backcrossing to C57BL/6J animals, a homozygous Kit Δ30k line was established.

All other mouse lines in this study were generated through cytoplasmic injection of mRNA Cas9 and gRNAs (Supplementary Table 2), as described in Korablev et al., 2019. For generation Pdgfra Δ2k and Pdgfra Δ60k lines CRISPR components were injected in C57BL/6J zygotes. For generation Kit Δ30k+ line CRISPR components were injected in Kit Δ30k zygotes.

All the procedures and technical manipulations with animals were in compliance with the European Communities Council Directive of 24 November 1986 (86/609/EEC) and approved by the Bioethical Committee at the Institute of Cytology and Genetics (Permission N45 from 16 November 2018).

### Cell cultures

#### Mouse embryonic fibroblasts (MEFs)

MEFs were obtained from E13.5 as described in^68^. MEFs were cultured at 37°C under 5% CO_2_ in Dulbecco’s Modified Eagle Medium (DMEM) (Thermo Fisher Scientific, 12800082), supplemented with 10% FBS (Capricorn Scientific, FBS-11A), 1x penicillin & streptomycin 10x (Capricorn Scientific, PS-B), 1x GlutaMax-I 100× (Thermo Fisher Scientific, 35050061). For subculture cells were rinsed with 1× PBS and detached using 0.25% trypsin-EDTA at 37°C for 3 min. Cells were typically split every 2–3 days at a 1:2 ratio. Flow cytometry showed more than 80% *Pdgfra* positive cells (Supplementary Fig. 2).

#### Melanocyte cell culture

Melanocytes from Wt and Kit Δ30k mice were obtained according to the Murphy et al., 2019 protocol^69^ with modifications. In brief, 4-day-old pups were decapitated, the skin was separated, rinsed in 10x penicillin/streptomycin solution (Capricorn Scientific, PS-B), and incubated overnight in 0.25% trypsin/DMEM (Capricorn Scientific, TRY-2B). Next day, blood, adipose and muscle tissues were removed, epidermal and dermal layers were disassociated. Epidermis was chopped with scissors in 0.25% trypsin/PBS solution, and additionally incubated in 0.25% trypsin for 10-15 minutes at 37°C. Then, the cell suspension was filtered through a nylon mesh to remove large tissue segments, cells were pelleted by centrifugation and pre-plated for 25-30 minutes. Finally, unattached cells were transferred to the plastic surface 6-well plate, covered with 0.1% Gelatin from porcine skin type B (Sigma, G9391), and cultivated in Melanocyte growth medium (PromoCell, C-39410), containing geneticin 100 ng/ml (G418) (Thermo Fisher Scientific, 11811023), for 48 hours. On day 3, the medium was replaced with a fresh one, without G418. At that step individual melanocytes could be clearly visualized. In 10-14 days, the majority of the cell population represented granulated melanocytes. The cells were grown for 4-7 weeks until the required cell quantity was reached and more than 80% of cells were *Kit* positive (Supplementary Fig.2).

#### Mast cell culture

Mast cells were differentiated from hematopoietic cells derived from bone marrow according to the Vukman et al., 2014 protocol^70^. Cells were cultured for 8 to 10 passages in Iscove’s Modified Dulbecco’s Medium (IMDM) (Thermo Fisher Scientific, 12200069) supplemented with 10% FBS (Thermo Fisher Scientific, 16141002), mrSCF(10 ng/ml) (Biolegend, 579702) and mrIL-3 (10 ng/ml) (Biolegend, 575502). Mast cells were also stained by toluidine blue (Biovitrum, 07-002) — a basic thiazine metachromatic dye that stains nuclei blue and has a high affinity for acidic granules, in accordance with the manufacturer’s recommendations.

According to the flow cytometry analysis of *Kit* and *FcεR1* expression, at the final passages mast cells accounted for ~90% of the total cell population (Supplementary Fig. 2). For the Hi-C experiment *Kit* positive cells were magnetically separated (Miltenyi Biotec, 130-097-146).

### Flow cytometry (FC)

FC was performed as described in^71^. A total of ~0.5-1 * 10^6^ cells/ml were harvested and washed in PBS. Antibody and cell suspension were mixed in an ice-cold FC buffer (PBS, 10% FBS) in a ratio 1:100 and incubated on ice for 1 hour. After incubation cells were pelleted by spinning at 300×g for 10 min. The pellet was washed in the FC buffer twice to remove unnecessary antibodies. The pellet was resuspended in a 500 μL buffer and filtered through nylon mesh to remove cell conglomerates. A negative control was prepared as a cell suspension washed in PBS without immunostaining. To exclude cells with compromised membranes cell suspension was incubated with 7-aminoactinomycin D (7-AAD) (Thermo Fisher Scientific, A1310) in a ratio 1:1000 for at least 20 min, after which the cell pellet was washed as described above. The prepared cell suspensions were analysed using BD FC Aria III (BD Biosciences).

### ChIP-seq

The anti-histone H3 (acetyl K27) ChIP-seq and input libraries were prepared for each cell type as described in Pekowska et al., 2018^72^ with slight modifications. The anti-CTCF ChIP-seq was performed in MEFs and mast cells. Briefly, ~ 5 * 10^6^ cells were harvested per experiment and crosslinked with 1% formaldehyde at RT for 10 min. Crosslinking was quenched with 125 mM glycine. Cells were lysed on ice in 0.5 ml lysis buffer (10 mM Tris pH 7.5, 1 mM EDTA, 0.4-1% SDS, 0.1% sodium deoxycholate, 1% Triton X-100, Complete Mini EDTA-free protease inhibitors (Roche, 11836170001)). Lysed chromatin was fragmented using the BANDELIN SONOPULSE until reaching a fragment size of 150–500 base pairs (5-10 cycles, 30s/60s on/off, 60% amplitude, 4°C). SDS concentration and sonication cycles were adjusted according to a cell type and shearing efficiency. Lysates were clarified by 16,000g centrifugation for 10-15 min at 4°C and diluted with 1-1.5 ml of lysis buffer without SDS to reduce SDS concentration before the immunoprecipitation step (~0.17% SDS final). Chromatin was pre-cleared with 20 μl Protein A magnetic beads (New England Biolabs, S1425S) for 1.5 h and 100 μL of the solution was saved as an input control. During this time, a 40 μl aliquot of Protein A beads was washed with PBS and rotated with the respective antibody at 4°C (1.5 - 5 μg antibody). Protein-DNA complexes were then immunoprecipitated overnight at 4°C with rotation. The beads were washed at 4°C in the following buffers: lysis buffer (0.1% SDS, twice), lysis buffer containing 0.5 M NaCl (twice), LiCl buffer (0.25 M LiCl, 0.5% IGEPAL-630, 0.5% sodium deoxycholate, twice), TE (pH 8.0) plus 0.2% Triton X-100 (once), and TE (pH 8.0, once). Crosslinks were reversed at 65°C overnight and DNA was purified with ChIP DNA Clean & Concentrator kit (Zymo Research, D5205). Sequencing libraries were prepared using the KAPA Hyper Prep kit. The anti-histone H3 (acetyl K27) (Abcam, ab4729) and anti-CTCF (Abcam, ab70303) antibodies have been successfully validated by Western Blot (Supplementary Fig.13). The experiments were performed in two biological replicates.

### Capture Hi-C (cHi-C)

Hi-C from two biological replicates of each cell type were performed as previously described in ^73^ with some modifications from^74^. A total of ~ 5 * 10^6^ cells were crosslinked in 1-2% PFA/PBS buffer at RT. A cell pellet was resuspended in a lysis buffer (150 mM Tris-HCl pH 7.5, 140 mM NaCl, 0.5% NP-40, 1% Triton X-100, 1x Complete Mini EDTA-free protease inhibitors) and incubated for 15 min at 4°C followed by 15 min at RT. Extracted nuclei were spun down at 800g then washed once in lysis buffer and once in NEBuffer3.1 (New England Biolabs, B703S). After centrifugation, nuclei pellet was resuspended in NEBuffer3.1 supplemented with 0.3% SDS, then incubated at 37°C for 1 h followed by 10 min at 65°C to open chromatin. 10% Triton X-100 was added to the reaction (1.8% final) to quench SDS and incubated at 37°C for 30 min. Chromatin was digested overnight with 400U DpnII (New England Biolabs, R0543M). Following the inactivation of DpnII, the nuclei were spun down at 2,500 g and resuspended in NEBuffer2.1. Biotin fill-in was performed with biotin-14-dATP (Invitrogen, 19524016) using Klenow DNA polymerase (SibEnzyme, E325) at 22°C for 4 h. Samples were centrifuged and the pellet was resuspended in ligation mixture (1xT4 Ligation buffer (New England Biolabs, B0202S), 5% PEG, 1% Triton X-100, 0.1 mg/ml BSA and T4 ligase). Ligation was carried out in 1.2 mL volume overnight at 16°C with mixing. The ligation products were reverse crosslinked at 65°C overnight with proteinase K (New England Biolabs, P8107S) and DNA was extracted with phenol-chloroform. Efficiency of DNA digestion and ligation was assessed with gel electrophoresis. Sonication was performed on the Covaris M220 system to obtain 200-400 bp fragments. DNA fragments were double size-selected using AMPure XP beads (Beckman Coulter, A63880). MyOne Streptavidin C1 magnetic beads (Invitrogen, 65001) were added to bind biotin-tagged fragments. Sequencing libraries were prepared using the KAPA Hyper Prep kit. The hybridization of libraries with RNA probes was performed according to the myBaits Manual v4.01 (Arbor Biosciences). Enrichment probes were designed over the region chr5:74,135,000-76,410,000.

### RNA-seq

0.2*10^6^–2*10^6^ cells were harvested and washed twice with PBS. Total RNA was extracted using Aurum Total RNA mini kit (Bio-Rad, 7326820). Sample concentration was evaluated using Qubit RNA HS (Q32852, Invitrogen). Gel electrophoresis was used to verify the integrity of extracted RNA. Three replicates were generated for each cell type and genotype, respectively.

### Immunohistochemistry (IHC)

E13.5-15.5 mice were fixed with 4% PFA solution in 1X PBS. Skin tissue sections from 4-day-old mice were harvested from head and back in 1X PBS and fixed in 4% PFA solution overnight on a roller shaker at 4°C. Next day samples were washed in 1X PBS solution three times for 30 min. For tissue dehydration organs were sequentially incubated for at least 24 h with a 15% and 30% sucrose solution in 1X PBS at 4°C. Next, organs were embedded in Tissue-Tek O.C.T. Compound (Sakura Finetek, 4583) and frozen. Organ sections of 50 um thickness were prepared on MICROM HM 505N cryostat (Microm) and immediately collected on SuperFrost™ slides (Thermo Fisher Scientific). Sections were washed with 1X PBS and incubated in blocking solution: 2% BSA (Sigma Aldrich, A2153), 0.2% Triton X-100 (Amresco, X100), 5% FBS (Capricom Scientific, FBS-11A). Primary antibodies anti-Kdr (R&D, AF644) and anti-Kit (CellSignaling, 3074) were diluted in blocking solution (1:100) and incubated overnight at slow speed on an orbital shaker at room temperature. Next, slices were washed with 1X PBS three times for 20 min and stained with secondary antibodies (Jackson Immuno Research, 705-165-147 and 711-545-152) and DAPI diluted in 1X PBS for 2 h at room temperature. Slices were washed with 1X PBS three times for 20 min and completely dried out and finally were mounted with ProLong™ Diamond Antifade Mountant (Thermo Fisher Scientific, P36965). Immunofluorescence were visualized under confocal fluorescence microscope LSM 780 NLO (Zeiss) with ZEN software (Zeiss). Microscopic analysis was carried out at the Multiple-access Center for Microscopy of Biological Subjects of the Institute of Cytology and Genetics SB RAS.

### Computational methods

All computations except the RNA-seq data analysis were performed using nodes of Novosibirsk State University high-throughput computational cluster.

#### cHi-C data processing

The cHi-C sequencing data were processed using Juicer software^75^ and mapped to the mm10 genome. Contact maps were generated from read pairs with MAPQ≥30 and normalized using the VC_SQRT method. Maps at 5 kb resolution were further visualized via the Epigenome browser^76^.

#### TAD organization analysis

The Hi-C maps for human (hg38), mice (mm10), rabbit (oryCun2), dog (canFam3), chicken (galGal5) were obtained from GSE167579^22^. The Hi-C map of african clawed frog (xenLae2) was obtained from^23^.

The conservativity of the *Kit* locus was estimated by the C-InterSecture software^21^. For the cross-species comparison, we generated maps of synteny regions from the pairwise genome alignments. At first, the genomes of interest were aligned by LastZ^77^. Then, the given alignments were converted into net-files by KentUtils [https://github.com/ucscGenomeBrowser/kent] and transformed into the syntenic maps by the C-InterSecture script. Additionally, with a considerable part of the genomes being not included in any synteny blocks, we mechanically filled gaps (< 1Mb) splitting codirected synteny blocks. The resulting synteny maps were used to liftover Hi-C contacts between the genomes by a “balanced” model.

#### Computation of the contacts enrichment

The comparison of capture Hi-C of wild type and mutant was performed by the C-InterSecture software, too. The synteny maps were manually generated according to the coordinates of deletion. Then, the Hi-C contacts of wild type were liftovered on the mutant genome by a “balanced” model, as though the deletion effects on chromosome architecture only by changing of a genomic distance. Finally, the given contacts were liftovered from mutant genome on wild type by an “easy” model accounting only synteny between locus.

The difference between the based and liftovered Hi-C map was calculated thus:

Let i, j and k - is genome coordinate in bins. Then:

*V_i,j_* is the contact value between bins i and j.
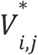 is the liftovered contact value between bins i and j.
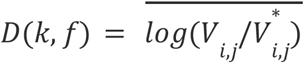 *for i* ∈ [*k* – *f,k*), *j* ∈ (*k, k* + *f*], *V_i,j_* ≠ 0, 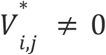, and *f* is distance in bins

Then, we performed a Z-score transformation of D(k,f) for all k and chosen f.

#### ChIP-seq data processing

Raw paired-end reads were preprocessed with Cutadapt tool to trim adapter sequences (-a and -A flags with Illumina adapter sequences)^78^. Read quality was assessed using FastQC. Coverage tracks (bigWig) were generated using Aquas pipeline with TF or histone parameters [https://github.com/kundajelab/chipseq_pipeline].

To determine a CTCF motif orientation we used the FIMO tool from MEME^79^ with the matrices MA0139.1 and MA1929.1 from JASPAR^80^.

#### RNA-seq data processing

Sequencing data were uploaded to and analyzed on the Galaxy web platform usegalaxy.org ^81^. Quality of sequencing data was controlled via FastQC Galaxy Version 0.73^82^. Reads were mapped to the mouse reference genome mm10 (GRCm38) with RNA STAR Galaxy Version 2.7.8a^83^ (default parameter) using the Gencode main annotation file release M1 (NCBIM37). Mapped reads were counted using featureCounts Galaxy Version 2.0.1^84^ (default parameter). Differential gene expression and normalized counts were calculated using DESeq2 Galaxy Version 2.11.40.7^85^ (default parameter).

To evaluate alleles activity in hybrid melanocytes we used *Mus castaneus* SNPs identified by the Mouse Genomes Project consortium^86,87^. For each gene, we calculated exonic SNPs coverage in the bam file and then calculated the average value across the replicates.

## Data availability

The raw sequencing data have been deposited in the NCBI SRA database with the following accession number PRJNA838252. Processed data, including Hi-C contact maps, RNA-seq and ChIP-seq tracks are available at https://genedev.bionet.nsc.ru/ftp/by_Project/Kit_locus_GEO.

## Contributions

N.B. conceived and supervised the study; E.K., A.R., V.L. and A.K. performed Hi-C, ChIP-seq, RNA-seq experiments; M.N. performed Hi-C data analysis; P.B. performed ChIP-seq and RNA-seq data analysis; T.S. performed immunohistochemistry analysis; A.K., G.K. and I.S. obtained genome–edited mice. All the authors contributed to the manuscript preparation.

## Acknowledgements

This work was supported with the budget project of Institute of Cytology and Genetics SB RAS (state project FWNR-2022-0019). Experiment with *Mus castaneus* hybrids was supported by Russian Science Foundation grant #22-14-00247. Hi-C experiments sequencing was performed using equipment of the Novosibirsk State University, supported by the Ministry of Education and Science of Russian Federation, grant #2019-0546 (FSUS-2020-0040). Illumina data analysis was supported by the strategic academic leadership program “Priority 2030” in Novosibirsk State University. Cell culturing was performed at the Collective Center of ICG SB RAS “Collection of Pluripotent Human and Mammalian Cell Cultures for Biological and Biomedical Research”, project number FWNR-2022-0019 (https://ckp.icgen.ru/cells/; http://www.biores.cytogen.ru/brc_cells/collections/ICG_SB_RAS_CELL). The calculations were performed using computational resources of Computational Center of Novosibirsk State University.

## References

1. Spielmann, M., Lupiáñez, D. G. & Mundlos, S. Structural variation in the 3D genome. NatRev Genet 19, 453–467 (2018).

2. Ibrahim, D. M. & Mundlos, S. Three-dimensional chromatin in disease: What holds us together and what drives us apart? Curr Opin Cell Biol 64, 1–9 (2020).

3. Hnisz, D. et al. Activation of proto-oncogenes by disruption of chromosome neighborhoods. Science 351, 1454–1458 (2016).

4. Lupiáñez, D. G. et al. Disruptions of topological chromatin domains cause pathogenic rewiring of gene-enhancer interactions. Cell 161, 1012–1025 (2015).

5. Redin, C. et al. The genomic landscape of balanced cytogenetic abnormalities associated with human congenital anomalies. Nat Genet 49, 36–45 (2017).

6. Kraft, K. et al. Serial genomic inversions induce tissue-specific architectural stripes, gene misexpression and congenital malformations. Nat Cell Biol 21, 305–310 (2019).

7. Cova, G. et al. Combinatorial effects on gene expression at the Lbx1/Fgf8 locus resolve Split-Hand/Foot Malformation type 3. 2022.02.09.479724 Preprint at https://doi.org/10.1101/2022.02.09.479724 (2022).

8. Laugsch, M. et al. Modeling the Pathological Long-Range Regulatory Effects of Human Structural Variation with Patient-Specific hiPSCs. Cell Stem Cell 24, 736–752.e12 (2019).

9. Guo, Y. A. et al. Mutation hotspots at CTCF binding sites coupled to chromosomal instability in gastrointestinal cancers. Nat Commun 9, 1520 (2018).

10. Flavahan, W. A. et al. Insulator dysfunction and oncogene activation in IDH mutant gliomas. Nature 529, 110–114 (2016).

11. Schwarzer, W. et al. Two independent modes of chromatin organization revealed by cohesin removal. Nature (2017) doi:10.1038/nature24281.

12. Nora, E. P. et al. Targeted Degradation of CTCF Decouples Local Insulation of Chromosome Domains from Genomic Compartmentalization. Cell 169, 930–944.e22 (2017).

13. Despang, A. et al. Functional dissection of the Sox9–Kcnj2 locus identifies nonessential and instructive roles of TAD architecture. Nature Genetics (2019) doi:10.1038/s41588-019-0466-z.

14. Williamson, I. et al. Developmentally regulated Shh expression is robust to TAD perturbations. Development (Cambridge) (2019) doi:10.1242/dev.179523.

15. Puputti, M. et al. Amplification of KIT, PDGFRA, VEGFR2, and EGFR in gliomas. Mol Cancer Res 4, 927–934 (2006).

16. Szerlip, N. J. et al. Intratumoral heterogeneity of receptor tyrosine kinases EGFR and PDGFRA amplification in glioblastoma defines subpopulations with distinct growth factor response. Proc Natl Acad Sci U S A 109, 3041–3046 (2012).

17. Battochio, A. et al. Detection of c-KIT and PDGFRA gene mutations in gastrointestinal stromal tumors: comparison of DHPLC and DNA sequencing methods using a single population-based cohort. Am J Clin Pathol 133, 149–155 (2010).

18. Benbarche, S. et al. Screening of ETO2-GLIS2-induced Super Enhancers identifies targetable cooperative dependencies in acute megakaryoblastic leukemia. Sci Adv 8, eabg9455 (2022).

19. Perrot-Applanat, M. & Di Benedetto, M. Autocrine functions of VEGF in breast tumor cells: adhesion, survival, migration and invasion. Cell Adh Migr 6, 547–553 (2012).

20. Grassot, J., Gouy, M., Perrière, G. & Mouchiroud, G. Origin and molecular evolution of receptor tyrosine kinases with immunoglobulin-like domains. Mol Biol Evol 23, 1232–1241 (2006).

21. Nuriddinov, M. & Fishman, V. C-InterSecture—a computational tool for interspecies comparison of genome architecture. Bioinformatics (2019) doi:10.1093/bioinformatics/btz415.

22. Li, D. et al. Comparative 3D genome architecture in vertebrates. BMC Biol 20, 99 (2022).

23. Hoencamp, C. et al. 3D genomics across the tree of life reveals condensin II as a determinant of architecture type. Science 372, 984–989 (2021).

24. Seifert, R. A., Alpers, C. E. & Bowen-Pope, D. F. Expression of platelet-derived growth factor and its receptors in the developing and adult mouse kidney. Kidney Int 54, 731–746 (1998).

25. Gouveia, L., Betsholtz, C. & Andrae, J. Expression analysis of platelet-derived growth factor receptor alpha and its ligands in the developing mouse lung. Physiol Rep 5, e13092 (2017).

26. Andrae, J., Gouveia, L., He, L. & Betsholtz, C. Characterization of platelet-derived growth factor-A expression in mouse tissues using a lacZ knock-in approach. PLoS One 9, e105477 (2014).

27. Millauer, B. et al. High affinity VEGF binding and developmental expression suggest Flk-1 as a major regulator of vasculogenesis and angiogenesis. Cell 72, 835–846 (1993).

28. Quinn, T. P., Peters, K. G., De Vries, C., Ferrara, N. & Williams, L. T. Fetal liver kinase 1 is a receptor for vascular endothelial growth factor and is selectively expressed in vascular endothelium. Proc Natl Acad Sci U S A 90, 7533–7537 (1993).

29. Rodríguez-Carballo, E. et al. The HoxD cluster is a dynamic and resilient TAD boundary *controlling the segregation of antagonistic regulatory landscapes*. Genes and Development (2017) doi:10.1101/gad.307769.117.

30. Narendra, V. et al. CTCF establishes discrete functional chromatin domains at the Hox clusters during differentiation. Science 347, 1017–1021 (2015).

31. Yoshida, H., Nishikawa, S.-I., Okamura, H., Sakakura, T. & Kusakabe, M. The Role of c-kit Proto-oncogene during Melanocyte Development in Mouse. In vivo Approach by the In utero Microinjection of Anti-c-kit Antibody. Development, Growth & Differentiation 35, 209–220 (1993).

32. Tacconi, C. et al. KIT is dispensable for physiological organ vascularisation in the embryo. Angiogenesis 25, 343–353 (2022).

33. Rajderkar, S. et al. Topologically Associating Domain Boundaries are Commonly Required for Normal Genome Function. 2021.05.06.443037 Preprint at https://doi.org/10.1101/2021.05.06.443037 (2021).

34. Rodríguez-Carballo, E. et al. Chromatin topology and the timing of enhancer function at the HoxD locus. Proc Natl Acad Sci U S A 117, 31231–31241 (2020).

35. Chahar, S. et al. >Context-dependent transcriptional remodeling of TADs during differentiation. 2022.07.01.498405 Preprint at https://doi.org/10.1101/2022.07.01.498405 (2022).

36. Banigan, E. J. et al. Transcription shapes 3D chromatin organization by interacting with loop-extruding cohesin complexes. 2022.01.07.475367 Preprint at https://doi.org/10.1101/2022.01.07.475367 (2022).

37. Zhang, S. et al. RNA polymerase II is required for spatial chromatin reorganization following exit from mitosis. Sci Adv 7, eabg8205 (2021).

38. Heinz, S. et al. Transcription Elongation Can Affect Genome 3D Structure. Cell 174, 1522–1536.e22 (2018).

39. Busslinger, G. A. et al. Cohesin is positioned in mammalian genomes by transcription,CTCF and Wapl. Nature 544, 503–507 (2017).

40. Zhang, S., Übelmesser, N., Barbieri, M. & Papantonis, A. Enhancer-promoter contact formation requires RNAPII and antagonizes loop extrusion. 2022.07.04.498738 Preprint at https://doi.org/10.1101/2022.07.04.498738 (2022).

41. Reed, K. S. M. et al. Temporal analysis suggests a reciprocal relationship between 3D chromatin structure and transcription. Cell Reports 41, (2022).

42. Olan, I. et al. Transcription-dependent cohesin repositioning rewires chromatin loops in cellular senescence. Nat Commun 11, 6049 (2020).

43. Hsieh, T.-H. S. et al. Resolving the 3D Landscape of Transcription-Linked Mammalian Chromatin Folding. Molecular Cell 78, 539–553.e8 (2020).

44. Barutcu, A. R., Maass, P. G., Lewandowski, J. P., Weiner, C. L. & Rinn, J. L. A TAD *boundary is preserved upon deletion of the CTCF-rich Firre locus*. Nature Communications (2018) doi:10.1038/s41467-018-03614-0.

45. Anania, C. et al. In vivo dissection of a clustered-CTCF domain boundary reveals developmental principles of regulatory insulation. Nat Genet 54, 1026–1036 (2022).

46. Islam, Z. et al. Active enhancers strengthen insulation by RNA-mediated CTCF binding at TAD boundaries. 2021.07.13.452118 Preprint at https://doi.org/10.1101/2021.07.13.452118 (2021).

47. Zuin, J. et al. Nonlinear control of transcription through enhancer-promoter interactions. Nature 604, 571–577 (2022).

48. Wehrle-Haller, B., Morrison-Graham, K. & Weston, J. A. Ectopic c-kit expression affects the fate of melanocyte precursors in Patch mutant embryos. Developmental Biology 177, 463–474 (1996).

49. Soriano, P. The PDGF alpha receptor is required for neural crest cell development and for normal patterning of the somites. Development 124, 2691–2700 (1997).

50. Stephenson, D. a et al. Platelet-derived growth factor receptor alpha-subunit gene (Pdgfra)*is deleted in the mouse patch (Ph) mutation*. Proceedings of the National Academy of Sciences of the United States of America 88, 6–10 (1991).

51. Korablev, A., Lukyanchikova, V., Serova, I. & Battulin, N. On-Target CRISPR/Cas9 *Activity Can Cause Undesigned Large Deletion in Mouse Zygotes*. International Journal of Molecular Sciences 21, 3604 (2020).

52. Greenwald, W. W. et al. Subtle changes in chromatin loop contact propensity are associated with differential gene regulation and expression. Nat Commun 10, 1054 (2019).

53. Gaspar-Maia, A., Alajem, A., Meshorer, E. & Ramalho-Santos, M. Open chromatin in pluripotency and reprogramming. Nat Rev Mol Cell Biol 12, 36–47 (2011).

54. Chabot, B., Stephenson, D. A., Chapman, V. M., Besmer, P. & Bernstein, A. The proto-oncogene c-kit encoding a transmembrane tyrosine kinase receptor maps to the mouse W locus. Nature 335, 88–89 (1988).

55. Geissler, E. N., Ryan, M. A. & Housman, D. E. The dominant-white spotting (W) locus of the mouse encodes the c-kit proto-oncogene. Cell 55, 185–192 (1988).

56. Nocka, K. et al. Expression of c-kit gene products in known cellular targets of W mutations in normal and W mutant mice--evidence for an impaired c-kit kinase in mutant mice. Genes & Development 3, 816–826 (1989).

57. Fontanesi, L. et al. The KIT Gene Is Associated with the English Spotting Coat Color Locus and Congenital Megacolon in Checkered Giant Rabbits (Oryctolagus cuniculus). PLoS ONE 9, e93750 (2014).

58. Hu, S. et al. KIT is involved in melanocyte proliferation, apoptosis and melanogenesis in the Rex Rabbit. PeerJ 8, e9402 (2020).

59. David, V. A. et al. Endogenous Retrovirus Insertion in the KIT Oncogene Determines White and White spotting in Domestic Cats. G3&amp;#58; Genes|Genomes|Genetics 4, 1881–1891 (2014).

60. Wong, A. K. et al. A de novo mutation in KIT causes white spotting in a subpopulation of German Shepherd dogs. Animal Genetics 44, 305–310 (2013).

61. Hauswirth, R. et al. Novel variants in the KIT and PAX3 genes in horses with white-spotted coat colour phenotypes. Animal Genetics 44, 763–765 (2013).

62. Huynh, L. & Hormozdiari, F. TAD fusion score: discovery and ranking the contribution of deletions to genome structure. Genome Biol 20, 60 (2019).

63. Real, F. M. et al. The mole genome reveals regulatory rearrangements associated with adaptive intersexuality. Science 370, 208–214 (2020).

64. Ringel, A. R. et al. Repression and 3D-restructuring resolves regulatory conflicts in evolutionarily rearranged genomes. Cell 185, 3689–3704.e21 (2022).

65. Battulin, N. et al. Comparison of the three-dimensional organization of sperm and fibroblast genomes using the Hi-C approach. Genome biology 16, 77 (2015).

66. Ryzhkova, A., Taskina, A., Khabarova, A., Fishman, V. & Battulin, N. Erythrocytes 3D genome organization in vertebrates. Scientific Reports (2021) doi:10.1038/s41598-021-83903-9.

67. Smirnov, A. V. et al. Evaluation of the α-casein (CSN1S1) locus as a potential target for a site-specific transgene integration. Scientific Reports 12, 1–10 (2022).

68. Kruglova, a a et al. Embryonic stem cell/fibroblast hybrid cells with near-tetraploid karyotype provide high yield of chimeras. Cell and tissue research 334, 371–80 (2008).

69. Murphy, B. M., Weiss, T. J. & Burd, C. E. Rapid Generation of Primary Murine Melanocyte and Fibroblast Cultures. J Vis Exp (2019) doi:10.3791/59468.

70. Vukman, K. V., Metz, M., Maurer, M. & O’Neill, S. M. Isolation and Culture of Bone Marrow-derived Mast Cells. Bio-protocol 4, e1053 (2014).

71. Flow cytometry (FACS) staining protocol (Cell surface staining). https://medicine.yale.edu/immuno/flowcore/protocols/analysis/.

72. Pękowska, A. et al. Gain of CTCF-Anchored Chromatin Loops Marks the Exit from Naive Pluripotency. Cell Systems 7, 482–495.e10 (2018).

73. Belaghzal, H., Dekker, J. & Gibcus, J. H. Hi-C 2.0: An optimized Hi-C procedure for high-resolution genome-wide mapping of chromosome conformation. Methods 123, 56–65 (2017).

74. Ulianov, S. V. et al. Active chromatin and transcription play a key role in chromosome partitioning into topologically associating domains. Genome Res 26, 70–84 (2016).

75. Durand, N. C. et al. Juicer Provides a One-Click System for Analyzing Loop-Resolution Hi-C Experiments. Cell Systems 3, 95–98 (2016).

76. Li, D. et al. WashU Epigenome Browser update 2022. Nucleic Acids Res 50, W774–781 (2022).

77. Harris, R. S. Improved pairwise alignment of genomic DNA. Ph.D. Thesis. (The Pennsylvania State University, 2007).

78. Martin, M. Cutadapt removes adapter sequences from high-throughput sequencing reads. EMBnet.journal 17, 10 (2011).

79. Bailey, T. L., Johnson, J., Grant, C. E. & Noble, W. S. The MEME Suite. Nucleic Acids Research 43, W39–W49 (2015).

80. Castro-Mondragon, J. A. et al. JASPAR 2022: the 9th release of the open-access database of transcription factor binding profiles. Nucleic Acids Research 50, D165–D173 (2022).

81. Afgan, E. et al. The Galaxy platform for accessible, reproducible and collaborative biomedical analyses: 2018 update. Nucleic Acids Res 46, W537–W544 (2018).

82. Babraham Bioinformatics - FastQC A Quality Control tool for High Throughput Sequence Data. https://www.bioinformatics.babraham.ac.uk/projects/fastqc/.

83. Dobin, A. et al. STAR: ultrafast universal RNA-seq aligner. Bioinformatics 29, 15–21 (2013).

84. Liao, Y., Smyth, G. K. & Shi, W. featureCounts: an efficient general purpose program for assigning sequence reads to genomic features. Bioinformatics 30, 923–930 (2014).

85. Love, M. I., Huber, W. & Anders, S. Moderated estimation of fold change and dispersion for RNA-seq data with DESeq2. Genome Biol 15, 550 (2014).

86. Keane, T. M. et al. Mouse genomic variation and its effect on phenotypes and gene regulation. Nature 477, 289–294 (2011).

87. Doran, A. G. et al. Deep genome sequencing and variation analysis of 13 inbred mouse strains defines candidate phenotypic alleles, private variation and homozygous truncating mutations. Genome Biol 17, 167 (2016).

