## Supplemental Files for "TAD border deletion at the *Kit* locus causes tissue-specific ectopic activation of a neighboring gene"

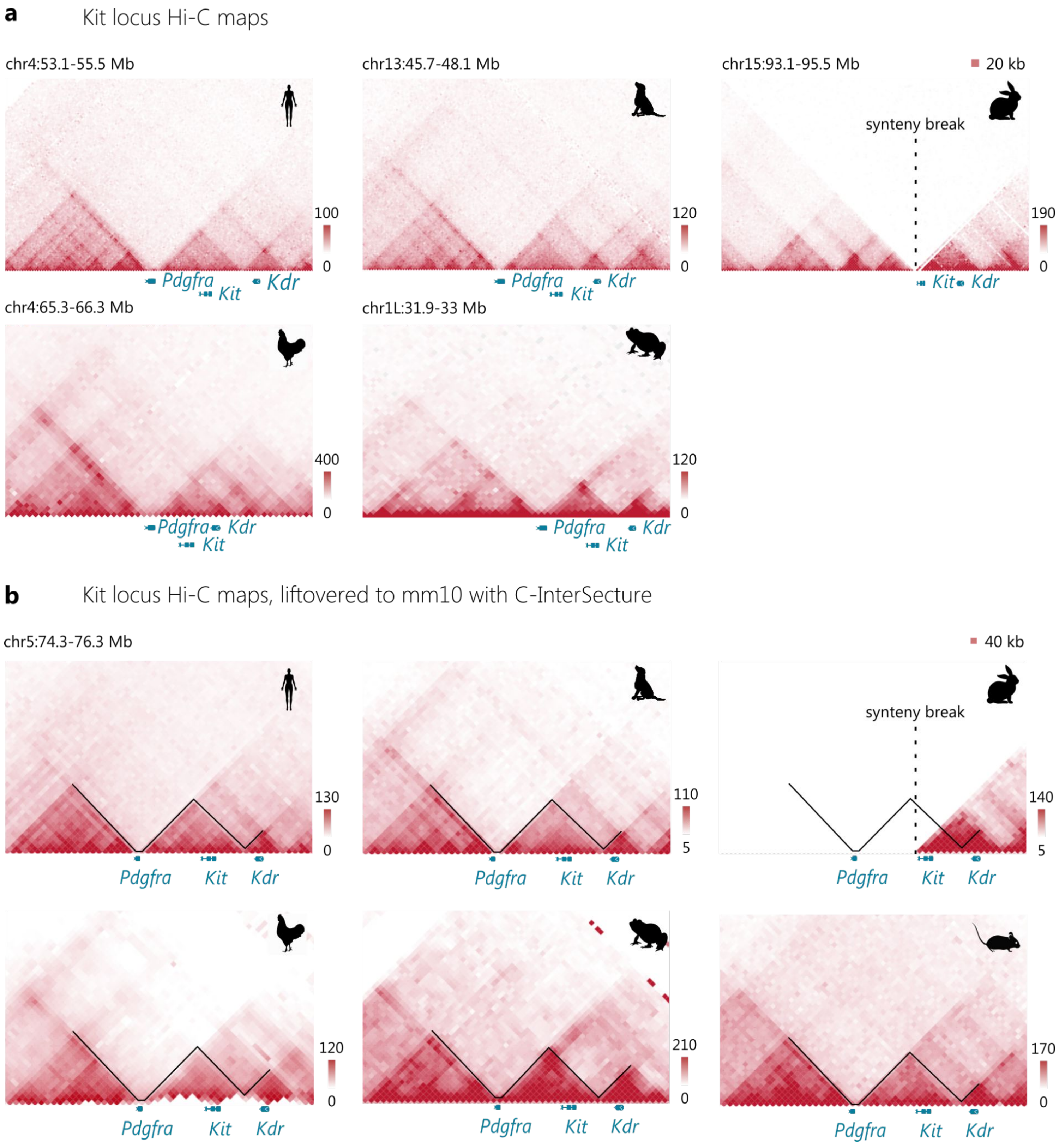


**Supplementary Figure 1. Hi-C heatmaps across six vertebrate species demonstrate conservativity of the TADs at the *Kit* locus. a** Hi-C contacts from human, dog, rabbit, chicken and frog. The Hi-C profile suggests that a synteny break between *Pdgfra* and *Kit* genes, observed in the rabbit data, may be due to a genome assembly error; **b** Hi-C contacts liftovered to the mouse genome. The C-InterSecture algorithm was used to neutralize the difference in the genomic distance between syntenic loci in different species. Colour bars reflect the interaction counts.

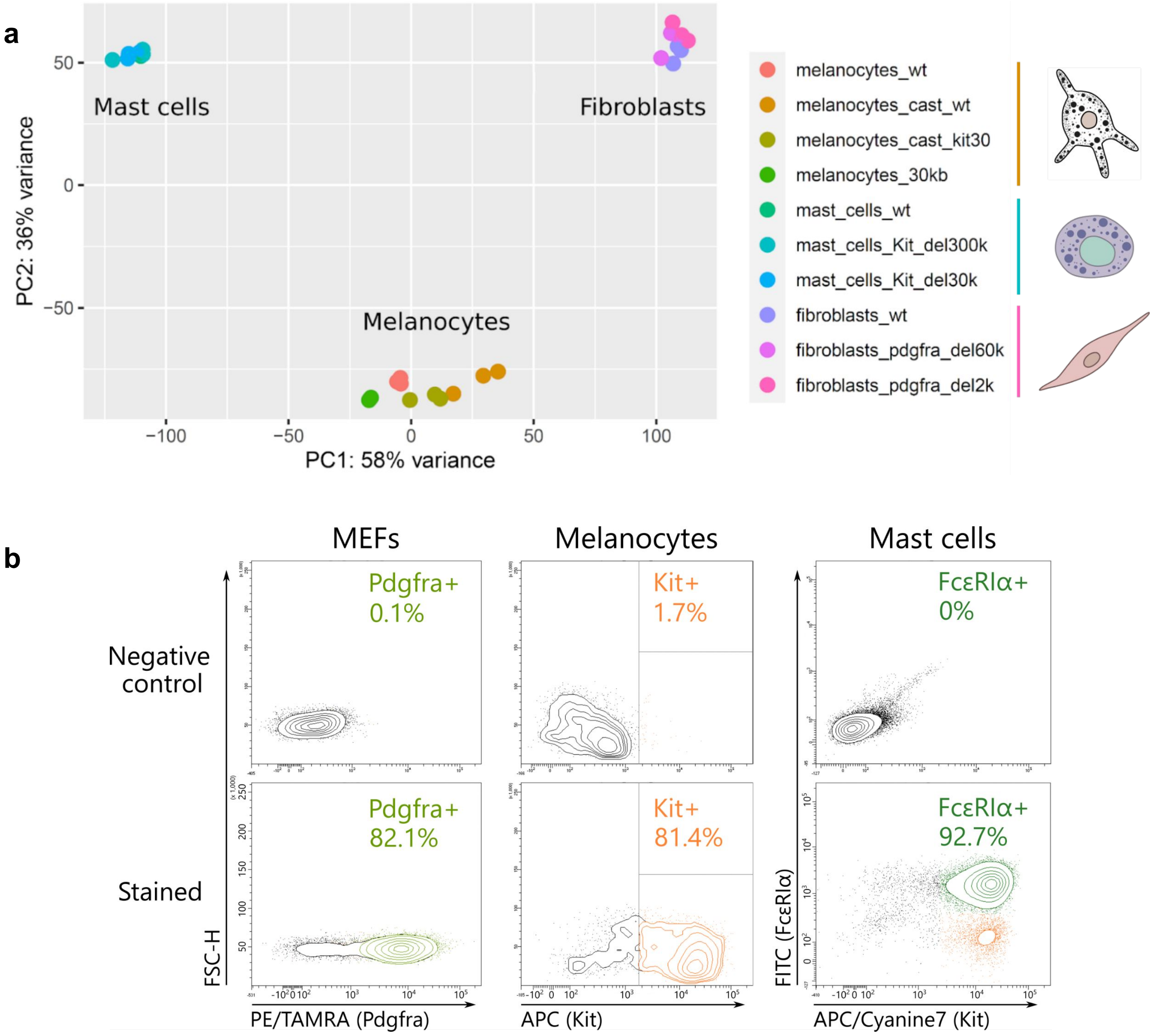


**Supplementary Figure 2. RNA-seq and flow cytometry analysis showing the purity of the obtained cell cultures. a** Plot of the principal component analysis indicates clearly distinguishable transcriptional profiles in the studied cell types and low variability between samples of the same cell type. Colored dots represent individual replicates; **b** Flow cytometry plots show average percentage of cell marker-positive cells in a population. Cells were stained with anti-Pdgfra (MEFs), anti-Kit (melanocytes), anti-Kit and anti-FcεRIα (mast cells) antibodies.

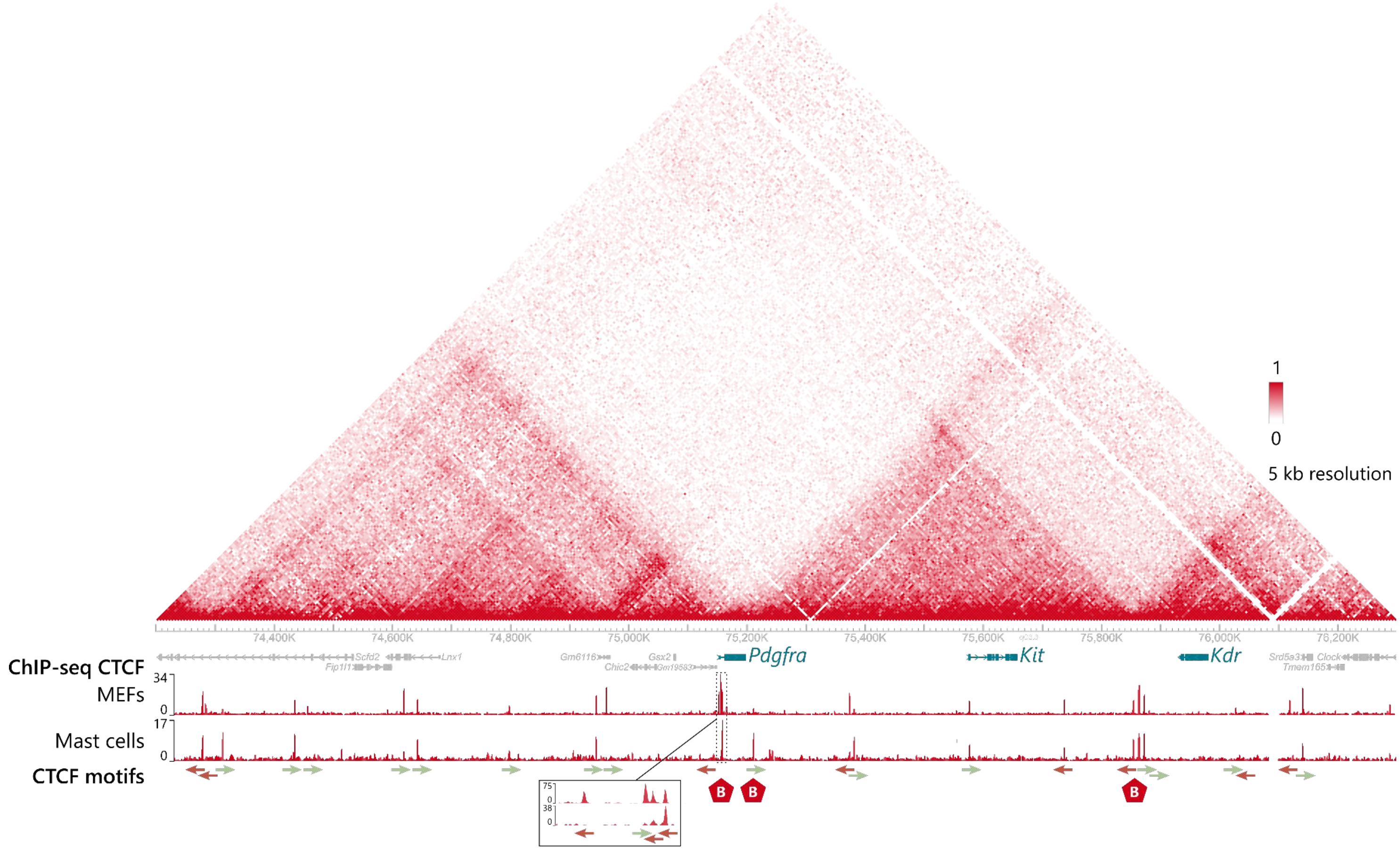


**Supplementary Figure 3. CTCF motif orientation at the *Pdgfra/Kit/Kdr* locus.** TAD borders are composed of at least two CTCF motifs at each border in convergent orientation (facing each other). A CTCF motif on plus DNA strand is forward oriented (right arrow), on minus DNA strand is reverse oriented (left arrow). Magnification of a 7 kb region shows a cluster of Pdgfra TAD boundary CTCF peaks and a Pdgfra promoter peak. Colour bar reflects the probability of chromatin contacts.

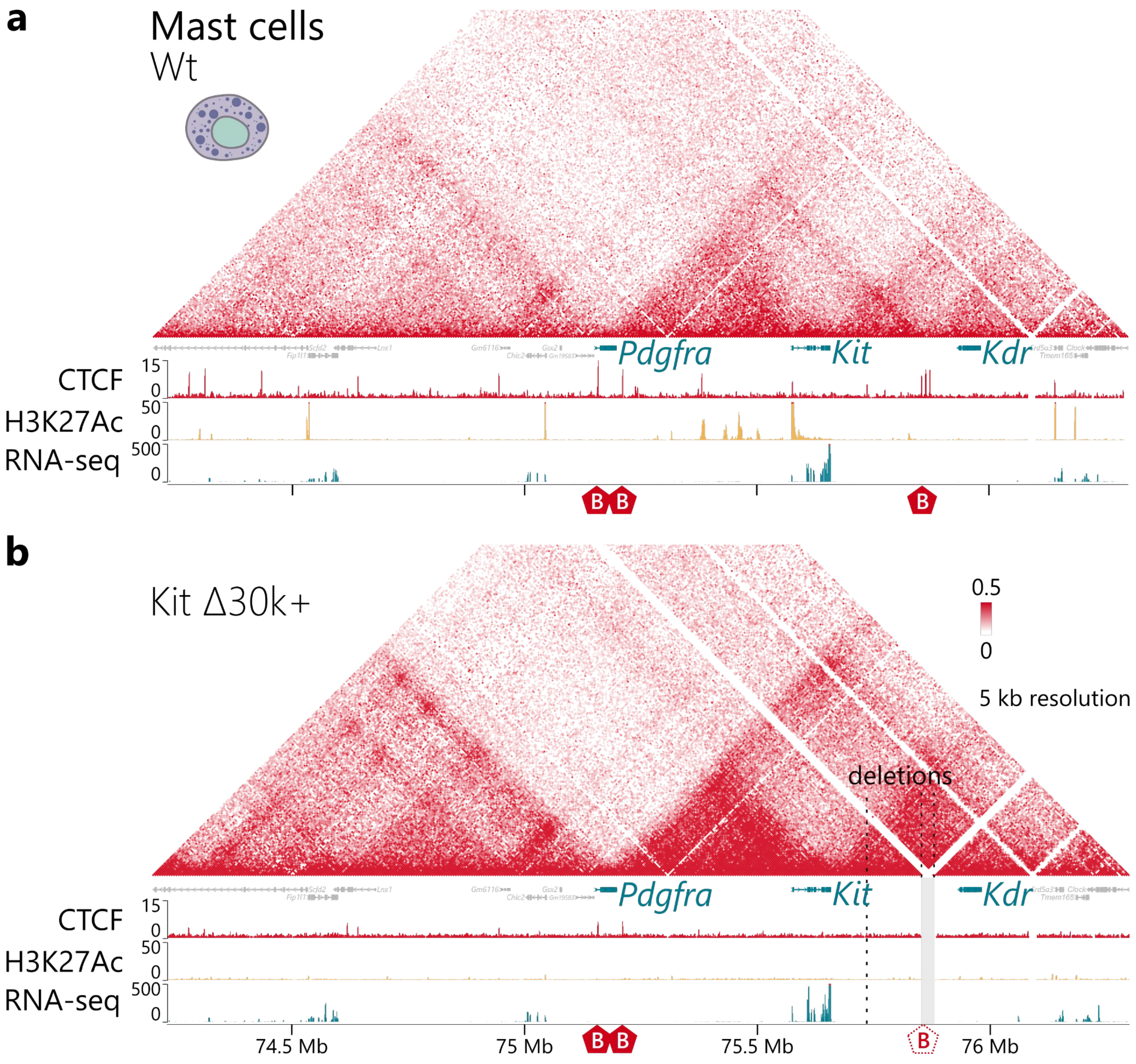


**Supplementary Figure 4. 3D genome organization of the *Kit* locus in mast cells.** Removal of boundary resulted in loss of inter-TAD insulation and extensive interactions across the TADs boundary. **a**-**b** cHi-C heatmaps, RNA-seq and ChIP-Seq signals across the *Kit* locus in wild-type (**a**) and KitΔ30k+ (**b**) mast cells. Colour bars reflect the interaction counts.

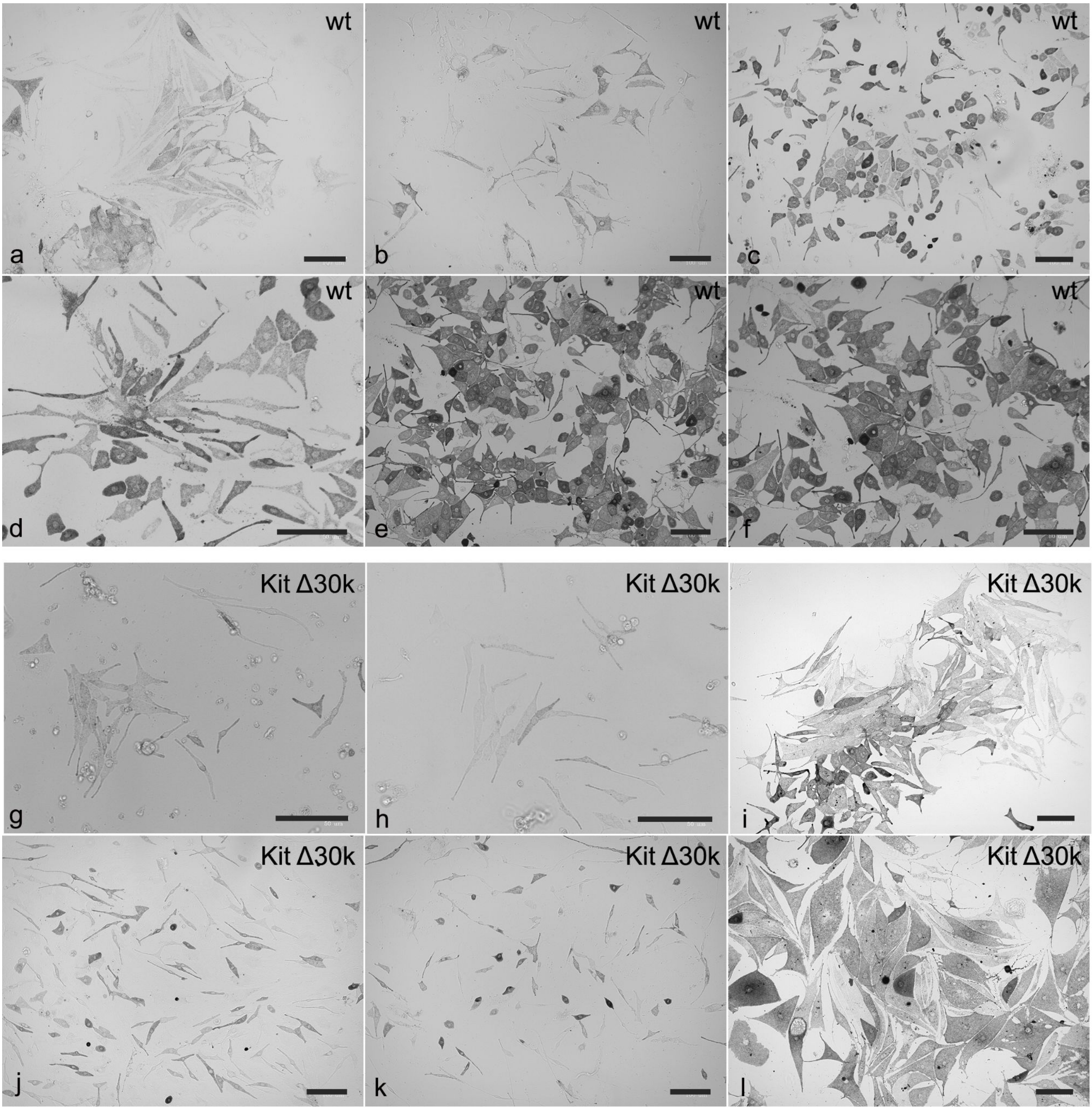


**Supplementary Figure 5. Melanocytes cell cultures at different culturing stages.** Bright-field images of differentiating wild-type (**a-f**) and Kit Δ30k (**g-l**) melanocytes. After 6-20 days in culture (**a-c** and **g-i**); after 21-30 days in culture (**d-f** and **g-l**). Scale bar: 100 um. A distinct feature of melanocytes morphology is visible: granules of dark pigment, melanin.

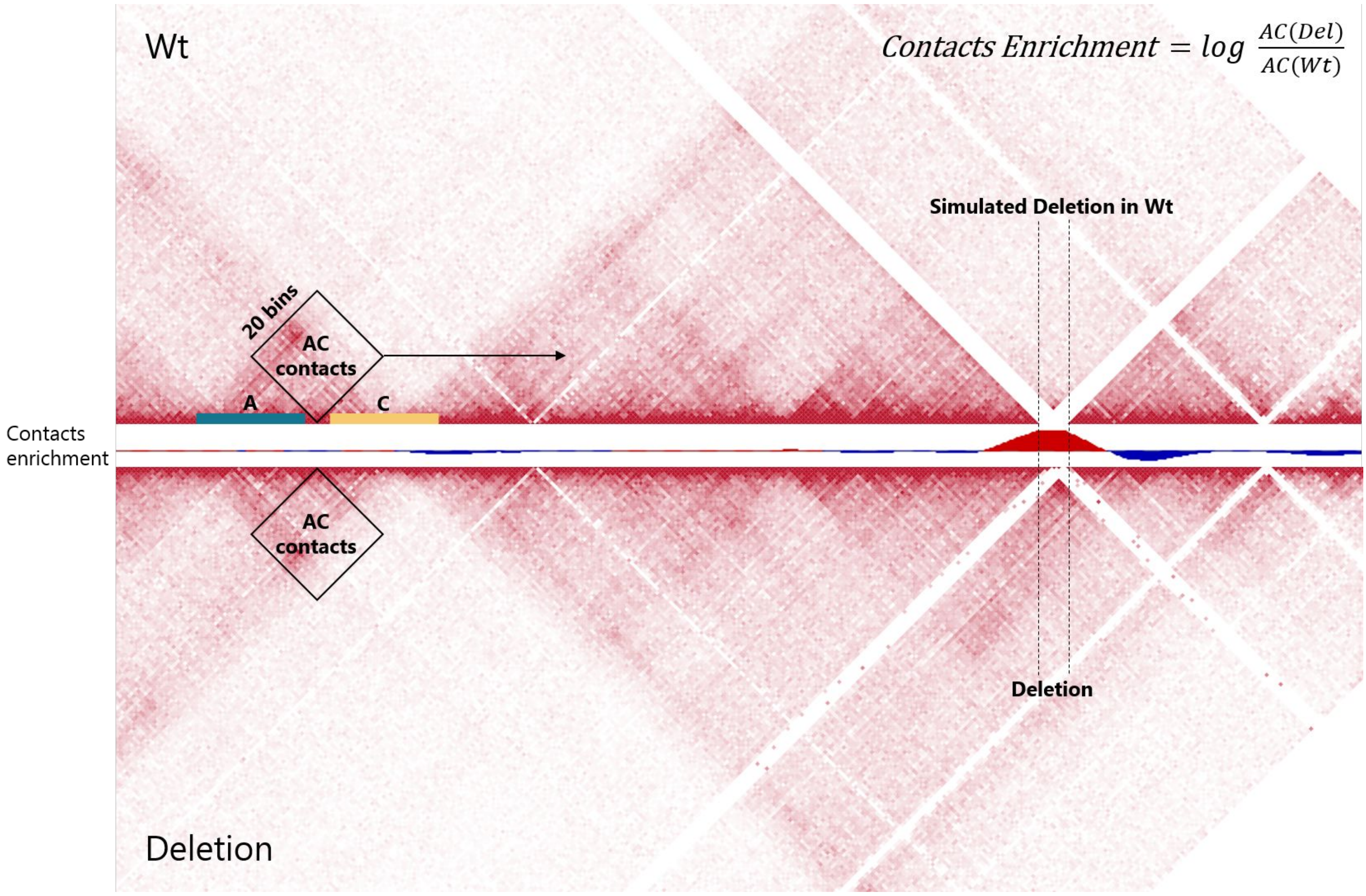


**Supplementary Figure 6. Schematic view of the contacts enrichment analysis in the mutant cHi-C data relative to the wild type.** Wild-type cHi-C reads were mapped to a custom genome containing a corresponding deletion. Matrices were VC_SQRT normalized. Then, log2-transformation of each Del contact / Wt contact ratio was performed. Next, we calculated the average contact ratio in a sliding window along the Hi-C diagonal. Obtained values were Z-score normalized. A series of different window sizes were tested (5–40 bins, with 1 bin = 5 kb). A 20 bins window size was used for the analysis.

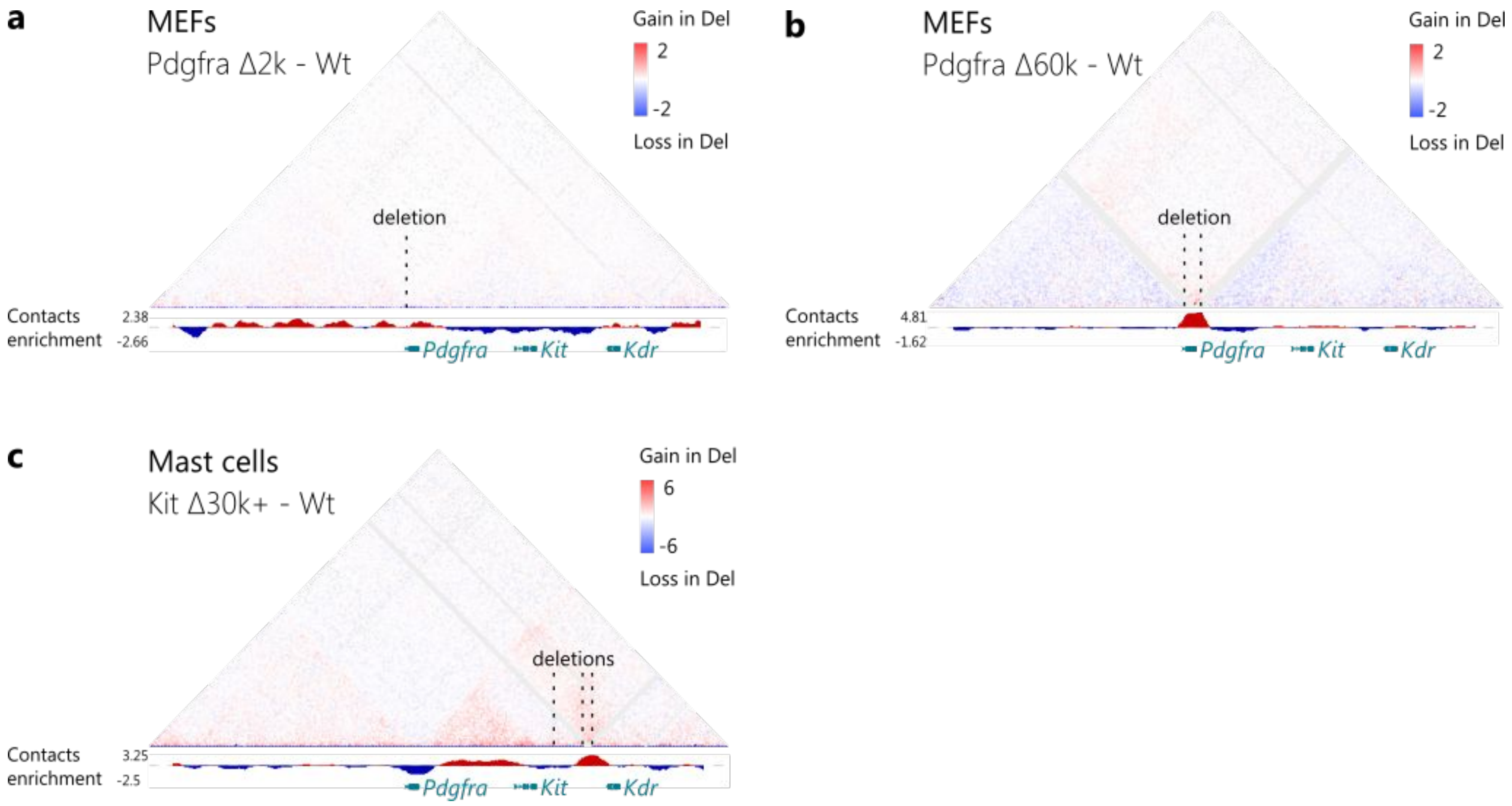


**Supplementary Figure 7. Impact of TAD boundary deletion on the inter-TADs contact frequency.** Subtraction maps show changes of contact frequencies in Pdgfra Δ2k (**a**) and Pdgfra Δ60k (**b**) MEFs, and KitΔ30k+ (**c**) mast cells, relative to the wild type. The tracks below the subtraction maps show the estimated contact enrichment, indicating loss of boundary insulation in Pdgfra Δ60k and KitΔ30k+ mutant cells, but not in Pdgfra Δ2k. Data is shown at 10 kb resolution.

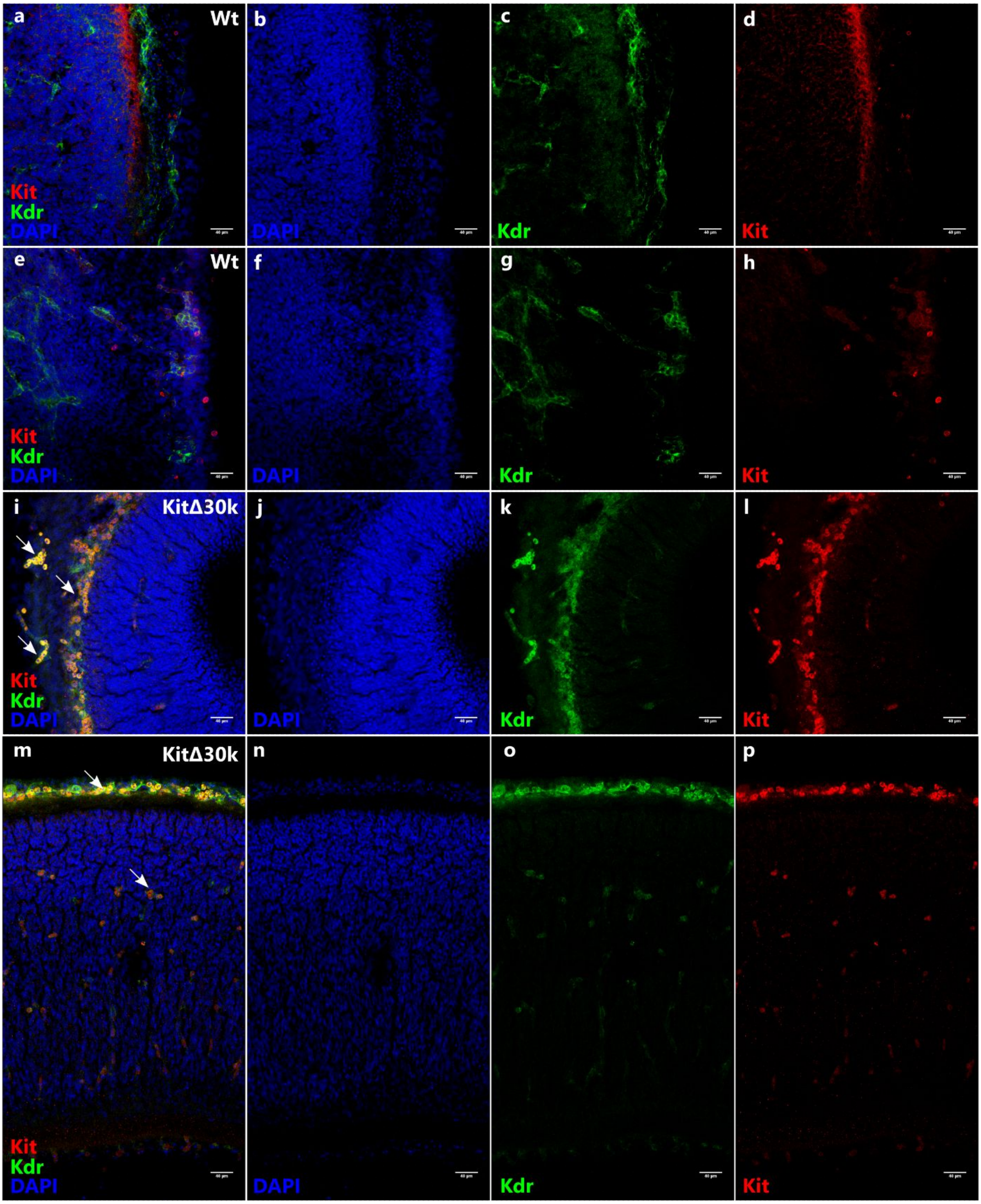


**Supplementary Figure 8. Detection of the *Kit-Kdr* double-positive cells in mouse embryonic skin.** Skin tissue sections from E13.5 wild-type (**a**-**h**) and Kit Δ30k (**j**-**q**) embryos. Images of representative regions of interest are shown: skin areas on the head (**a**-**d**, **i**-**l**) and on the back (**e**-**h**, **m**-**p**). Arrows: double-positive cells (Kit+Kdr+), located along the branching blood vessels in Kit Δ30k embryos. Scale bar: 40 um.

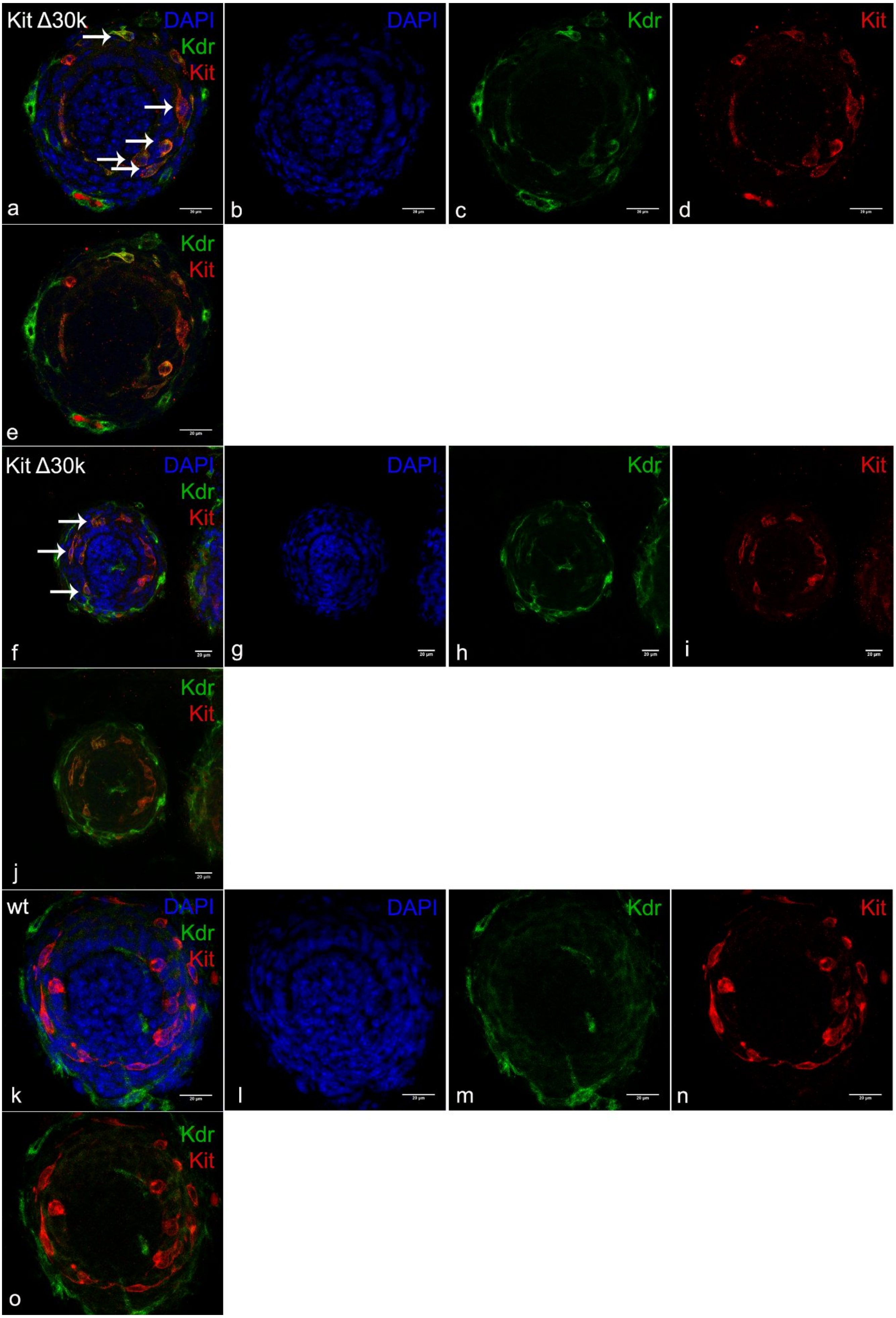


**Supplementary Figure 9. Detection of *Kit-Kdr* double-positive cells in vibrissae hair bulge of mouse embryo.** Representative vibrissae hair bulge sections of E15.5 embryos. **a-j** Kit Δ30k embryo. Double-positive cells (Kit+Kdr+) are indicated by arrows; **k-o** wild-type embryo. Scale bar 20 um

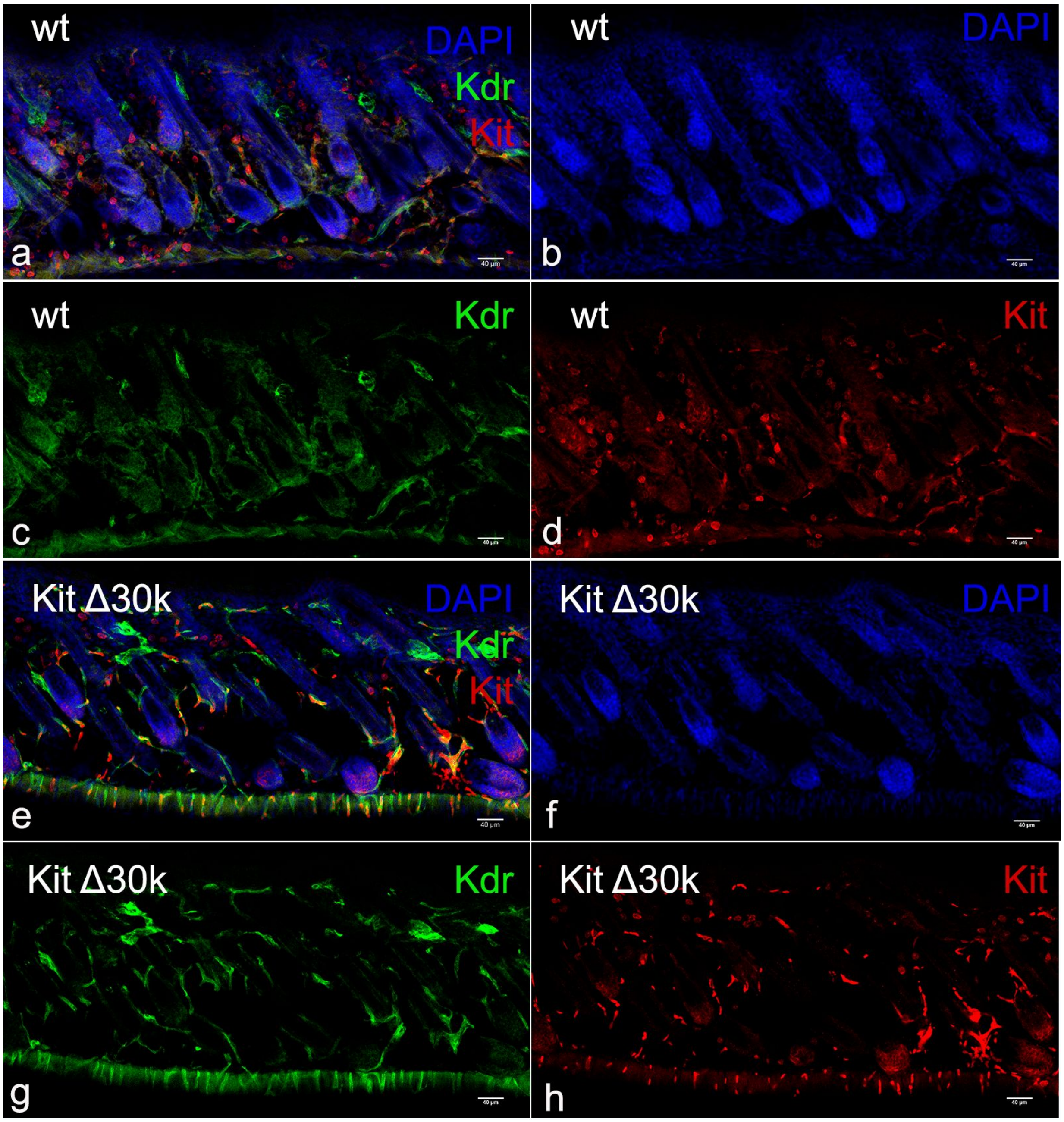


**Supplementary Figure 10. Detection of *Kit*-positive cells in the skin of 4-day-old pups.** Ratio of *Kit*-positive cells (red) localized in hair follicles and near them to all DAPI-stained nuclei inside the area was counted using ImageJ Cell Counter plugin. Six areas from different sections were analysed for each genotype. (**a-d**) wild-type, (**e-h**) Kit Δ30k. Scale bar: 40 um.

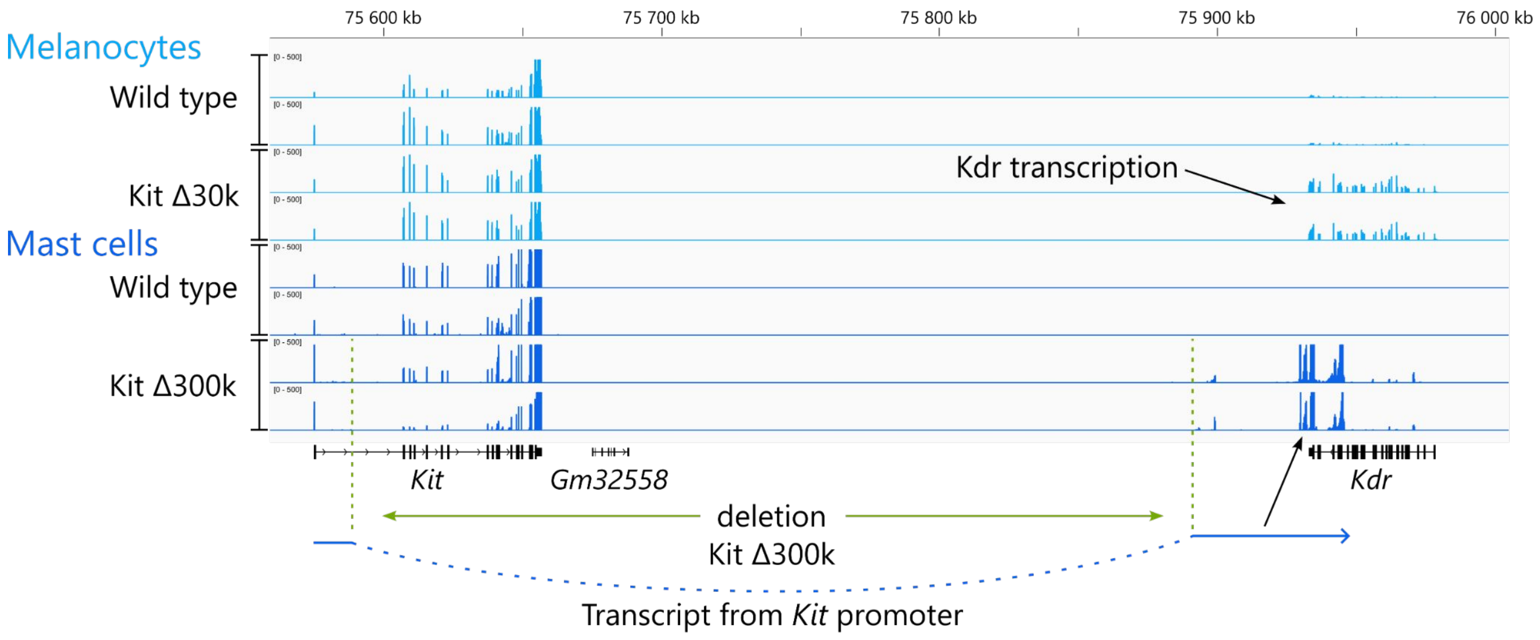


**Supplementary Figure 11. Expression pattern of the *Kit/Kdr* locus upon deletions of different size at the TAD border between Kit and Kdr TADs.** Mast cells from mice, carrying a 300 kb deletion at the border between Kit and Kdr TADs (Kit Δ300k), demonstrate appearance of an antisense *Kdr* transcript, as the transcript does not align with exons of *Kdr*. Whereas melanocytes of mice, carrying a 30 kb deletion at the same border (Kit Δ30k), demonstrate activation of *Kdr* transcription. The antisense transcription in Kit Δ300k could be explained by transcription started from the *Kit* promoter, which remained intact after the deletion, and continued to the closest termination site. Since *Kdr* locates on an opposing DNA strand to *Kit*, its transcript was antisense. The Kit Δ300k mice strain is heterozygous, therefore *Kit* was expressed from a normal allele.

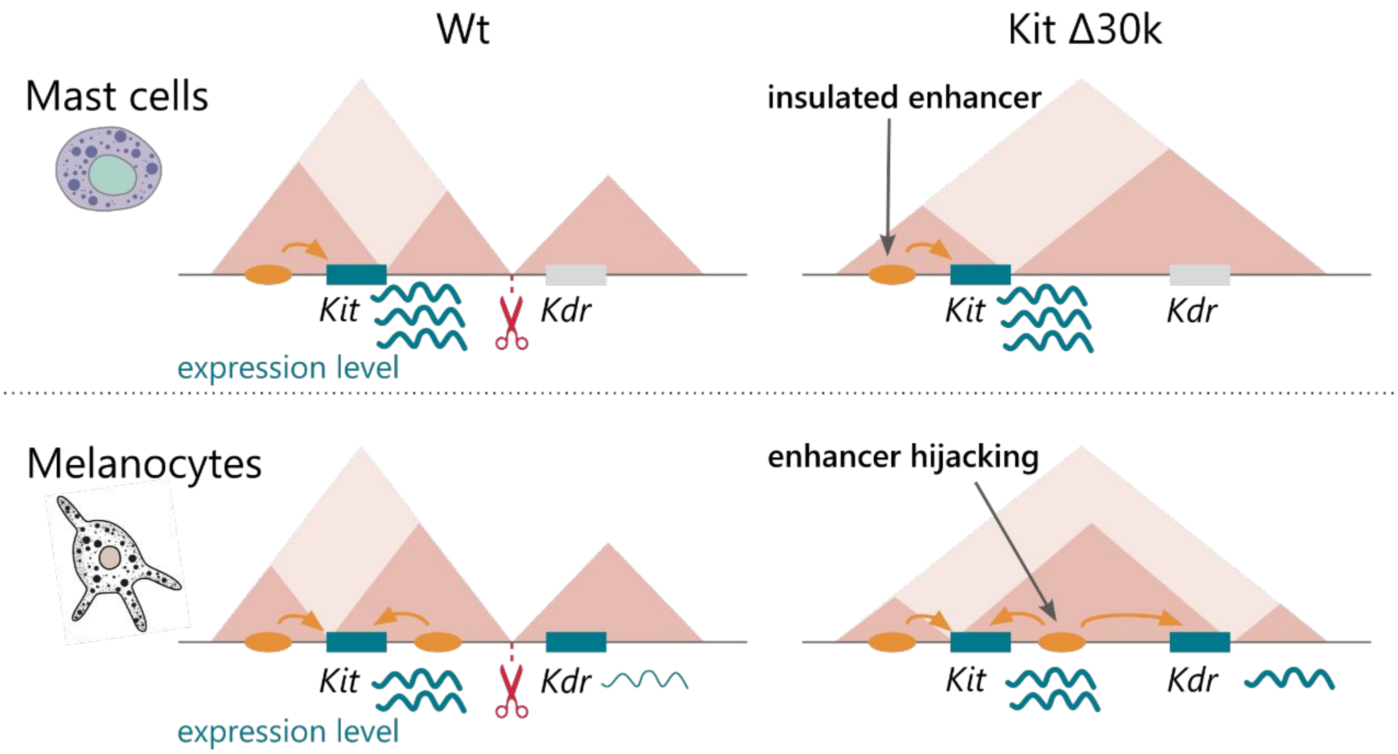


**Supplementary Figure 12. Schematic interpretation of the study results.** Deletion of CTCF binding sites at the border between Kit and Kdr TADs caused TADs fusion both in mast cells and melanocytes. Though enhancer hijacking and activation of *Kdr* transcription occured only in the mutant melanocytes.

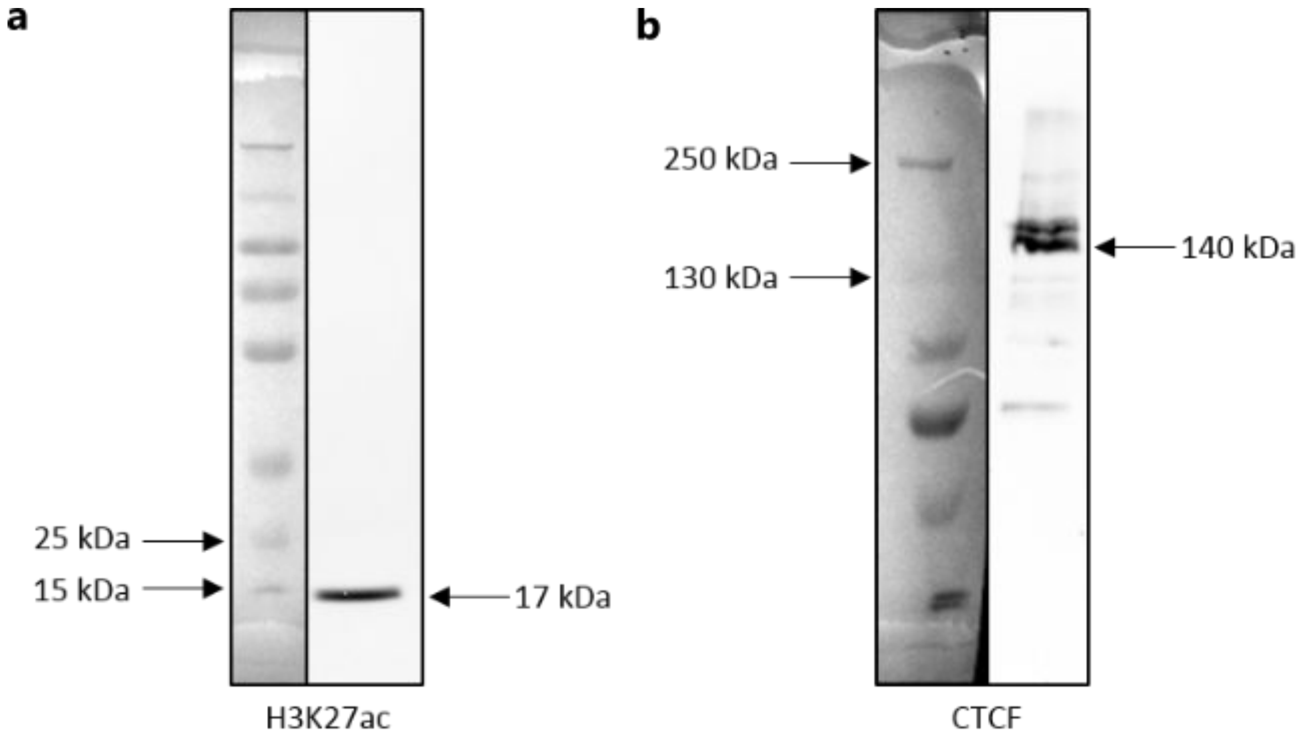


**Supplementary Figure 13. Western blot analysis for the antibodies validation.** Anti-H3K27ac (**a**) and anti-CTCF (**b**) were diluted 1:3000 for the Western blotting. mESC whole protein samples were used for the western blotting.

**Supplementary Table 1. Non-exhaustive list of studies concerning deletions of CTCF binding sites at TADs borders.**

| Locus | Model | Deletion size | Expressio n change | Reference |
|---|---|---|---|---|
| *Tbx5/Lhx5, Twist1/Ahr* | *In vivo* mouse | 21, 13 kb | Yes | Rajderkar S. et al., 2021 (bioRxiv) |
| *Smad7/Smad2, Sim1/Rfx6, Tbx3/Tbx5, Neurog2/Pitx2, Dmrt1,3/Dmrt2* | *In vivo* mouse | 11–72 kb | No | Rajderkar S. et al., 2021 (bioRxiv) |
| *Sox9/Kcnj2* | *In vivo* mouse | 6 kb | No* | Despang et al., 2019 |
| *HoxD* | *In vivo* mouse | 26 bp, 1.5 kb | Yes* | Rodríguez-Carballo et al., 2020 |
| *HoxD* | *In vivo* mouse | ~40–90 kb, 350 kb | Yes | Rodríguez-Carballo et al., 2017 |
| *HoxD* | *In vivo* mouse | ~20–30 kb | No | Rodríguez-Carballo et al., 2017 |
| *Epha4/Pax3* | *In vivo* mouse | 1.7 Mb | Yes | Lupiáñez et al., 2015 |
| *Shh/Mnx1* | mESCs | 35 kb | Yes | Williamson et al., 2019 |
| *Shh/Mnx1* | mESCs | 1 kb | No | Williamson et al., 2019 |
| *Tsix/Xist* | mESCs | 58 kb | Yes** | Nora et al., 2012 |
| *TAL1/CMPK1; LMO2/CAPRIN1* | HEK-293T | 400 bp; 25 kb | Yes | Hnisz et al., 2016 |
| *HoxA* | ESCs, motor neurons | 9 bp | Yes | Narendra et al., 2015 |
| *PDGFRA* | IDH wild-type glioma spheres | 10 bp | Yes | Flavahan et al., 2015 |

* moderate change
** long-range transcriptional misregulation

**Supplementary Table 2. List of Guide RNAs and primer sequences for mouse genotyping, used in the study.**

| gRNA | | |
|---|---|---|
| **Deletion** | **Protospacer sequence+PAM** | **Mm10 genomic coordinates** |
| Kit Δ30k | GCGCCGTAAGTGCTGAAAAGAGG<br>GAGGTAAGAGCAATCCGGTAAGG | chr5:75,852,777-75,852,799<br>chr5:75,881,199-75,881,221 |
| Kit Δ30k+ | ACAATACAAAATAATCTGGGGGG<br>AAAGCACTATGCCCAGTGAAAGG | chr5:75,736,367-75,736,389<br>chr5:75,736,660-75,736,682 |
| Pdgfra Δ2k | GTGGACCACCAATACTAGCTGGG<br>CAGTGGTTAAGGTCACCGAATGG | chr5:75,155,567-75,155,589<br>chr5:75,157,744-75,157,766 |
| Pdgfra Δ60k | GTGGACCACCAATACTAGCTGGG<br>GTCACCAAGTATGCGGTCATTGG | chr5:75,155,567-75,155,589<br>chr5:75,216,592-75,216,614 |

**Genotyping primers**

**For Pdgfra/Kit deletions (FWD-278+REV-279; FWD-278+REV-281; FWD-278+REV-308)**

| Name | Sequence 5'-3' |
|---|---|
| FWD-278 | CTGATTCCCGACCTCATCGG |
| REV-279 | CGTTTGCGACTCGTATCTCG |
| REV-281 | GGACTTTCCTTCCCTCGTCC |
| REV-308 | CTTATGGGCCTCTCGACTCG |

**For Kit/Kdr deletions (88+89; 90+93)**

| | |
|---|---|
| 88 | CCTACGAGCCTTCACGTTGT |
| 89 | TGAGGACCGCTGATAGGGAA |
| 90 | AAGGCTGTTGTACTGCGTGA |
| 93 | GTGCTATGGGAGCCGAAAGA |

### Supplementary References


1. Rajderkar, S. *et al.* Topologically Associating Domain Boundaries are Commonly Required for Normal Genome Function. 2021.05.06.443037 Preprint at https://doi.org/10.1101/2021.05.06.443037 (2021).

2. Despang, A. *et al.* Functional dissection of the Sox9–Kcnj2 locus identifies nonessential and instructive roles of TAD architecture. *Nat Genet* **51**, 1263–1271 (2019).

3. Rodríguez-Carballo, E. *et al.* Chromatin topology and the timing of enhancer function at the HoxD locus. *Proceedings of the National Academy of Sciences* **117**, 31231–31241 (2020).

4. The HoxD cluster is a dynamic and resilient TAD boundary controlling the segregation of antagonistic regulatory landscapes. http://genesdev.cshlp.org/content/31/22/2264.full.

5. Lupiáñez, D. G. *et al.* Disruptions of Topological Chromatin Domains Cause Pathogenic Rewiring of Gene-Enhancer Interactions. *Cell* **161**, 1012–1025 (2015).

6. Williamson, I. *et al.* Developmentally regulated Shh expression is robust to TAD perturbations. *Development* **146**, dev179523 (2019).

7. Nora, E. P. *et al.* Spatial partitioning of the regulatory landscape of the X-inactivation centre. *Nature* **485**, 381–385 (2012).

8. Activation of proto-oncogenes by disruption of chromosome neighborhoods | Science. https://www.science.org/doi/full/10.1126/science.aad9024.

9. Narendra, V. *et al.* CTCF establishes discrete functional chromatin domains at the Hox clusters during differentiation. *Science* **347**, 1017–1021 (2015).

10. Flavahan, W. A. *et al.* Insulator dysfunction and oncogene activation in IDH mutant gliomas. *Nature* **529**, 110–114 (2016).
